# Astrocyte Store-Released Calcium Modulates Visual Cortex Synapse Development and Circuit Function

**DOI:** 10.1101/2025.07.20.665758

**Authors:** Gillian Imrie, Jordan Mar, Madison Gray, Isabella Farhy-Tselnicker

**Affiliations:** Department of Biology, Texas A&M University, College Station, TX 77843, USA; Texas A&M Institute for Neuroscience (TAMIN) Texas A&M University, College Station, TX 77843, USA; Center for Biological Clocks Research Texas A&M University, College Station, TX 77843, USA

**Keywords:** Astrocyte, calcium, IP3R2, visual circuit development, synapse development

## Abstract

Astrocytes, a major class of glial cells, are critical regulators of synapse development during early postnatal life. While dysregulation of this process is implicated in numerous neurological disorders, the precise mechanisms by which astrocytes guide synapse formation and maturation remain poorly understood. A central signaling pathway for astrocytes is the dynamic fluctuation of intracellular calcium (Ca^2+^), which can arise from various sources and modulate a wide range of downstream effects. A key astrocytic mechanism for integrating neuronal signals is the release of Ca^2+^ from endoplasmic reticulum stores mediated by the IP3 Receptor Type 2 (IP3R2). Although defects in this signaling pathway have been mainly linked to adult brain dysfunction, its role in shaping synaptic development, a period when astrocyte-neuronal communication is established, is largely unknown. Here, we investigated the role of IP3R2-mediated Ca^2+^ signaling in astrocyte-dependent regulation of synapse development in the mouse visual cortex. Using a combination of histological, molecular, and circuit-level approaches, we found that loss of astrocytic IP3R2 leads to significant deficits in the maturation of glutamatergic but not GABAergic synapses. These synaptic disruptions were accompanied by attenuated visually evoked neuronal activation and impaired behavioral responses to visual threat stimuli. We further show that astrocytic morphological complexity is diminished in the absence of IP3R2, suggesting that store-released Ca^2+^ is required for both the structural and functional maturation of astrocyte-neuron interactions. Our findings establish a critical role for astrocytic IP3R2-mediated Ca^2+^ signaling in shaping excitatory circuit development and the emergence of visually driven behaviors.

## INTRODUCTION

Synapses, the fundamental units of electrochemical communication in the brain, are tightly regulated by astrocyte activity throughout all life stages^1–3^. During early postnatal development, when excitatory and inhibitory neuronal circuits are actively forming and refining, astrocytic regulatory functions influence the maturation, strength, and specificity of synaptic connections^2,4,5^. Dysregulation of synapse formation is linked to numerous brain pathologies, including depression and mood disorders^6^, autism spectrum disorders^7^, and epilepsy^8^; however, the precise mechanisms by which astrocytes regulate synapse formation and maturation are not yet fully understood.

In the developing mouse visual cortex (VC), synapse maturation follows a well-defined postnatal trajectory with onset of synaptogenesis occurring at postnatal day (P) 7, peaking at P14 around the period when the eyes open, and stabilizing near P28^2^. Astrocyte development is temporally aligned with these synaptogenic stages^2,9,10^. During these timepoints, astrocytes engage in robust bidirectional communication with neurons and undergo major changes in gene expression and structural morphology which promote the appropriate spatiotemporal recruitment of neuronal synaptic components in a neuronal-activity dependent manner^4,11^. These developmentally mediated transcriptomic changes allow astrocytes to dynamically signal in a rapidly changing environment and integrate neuronal activity in ways appropriate to developmental stage and context^4,11,12^. Though numerous reports describe the importance of these developmental interactions^2–5,13^, the exact signaling pathways astrocytes engage to modulate them remain unresolved.

A major mechanism underlying astrocytic responses to intrinsic and extrinsic cues occurs via fluctuations in intracellular calcium (Ca^2+^), which arise from diverse sources and can vary widely in spatial and temporal dynamics, and have been linked to multiple astrocytic regulatory functions from the single cell to behavioral level^14–17^. These include processes such as energy metabolism, neuronal circuit synchronization, network scaling, and encoding of important sensory stimulus details like salience and context^18–23^. A central pathway by which astrocytes integrate neuronal signals is through releasing stored Ca^2+^ from the endoplasmic reticulum (ER). Neurotransmitters and neuromodulators bind to astrocytic G-protein coupled receptors (GPCRs), activating phospholipase C to generate inositol 1,4,5-trisphosphate (IP3). IP3 binds to its receptors on the ER, which in cortical astrocytes occurs predominantly through IP3 Receptor Type 2 (IP3R2), triggering Ca^2+^ release into the cytosol which modulates a wide range of downstream effects^12,24–31^. While it has been shown that disruptions in IP3R2 mediated Ca^2+^ signaling are linked to some functional deficits in adults (including altered connectivity, cortical circuit dysregulation, and autism spectrum disorder like behavioral phenotypes^4,17,25,31,32^), little is known about the involvement of this signaling pathway in modulating synaptic development, a time when astrocyte-neuronal communication is established.

Here, we investigated the role of IP3R2 mediated store-released Ca^2+^ signaling in the astrocyte dependent regulation of synapse development of the mouse visual cortex. Using a combination of histological, molecular, and circuit level approaches across defined postnatal timepoints, we found that loss of astrocytic IP3R2 leads to deficits in the maturation of glutamatergic but not GABAergic synapses. These synaptic disruptions were accompanied by attenuated visually evoked neuronal activation and impaired behavioral responses to visual threat stimuli. We further show that astrocyte morphological complexity is diminished in the absence of IP3R2 at P16, a timepoint critical for astrocytic synapse regulation, suggesting that store-released Ca^2+^ is required for both the structural and functional maturation of astrocyte-neuron interactions. Our findings establish a critical role for astrocytic IP3R2-mediated Ca^2+^ signaling in shaping the development of excitatory circuits and visually driven behaviors.

## METHODS

### ANIMALS

All animal work was approved by the Texas A&M University Institutional Animal Care and use Committee (IACUC).

Mice were maintained by the Texas A&M University Comparative Medicine Program under standard housing conditions on a 12-hour light:dark cycle with *ad libitum* access to food and water. Both female and male mice from newborn through adult developmental timepoints were used in experiments. The following mouse lines were used: Wild-type (WT; C57Bl6/J) were purchased from Jackson Labs and bred in-house (Jax #000664). Mice were used for breeding and backcrossing, and as controls. IP3R2 KO (Itpr2^tm1.1Chen^) was originally obtained from the Ju Chen lab at UCSD^33^ and maintained on C57BL6/J background as KO X KO or het X het breeding schemes. To generate experimental groups, KO mice were compared with WT collected at the same developmental times.

## MOUSE TISSUE COLLECTION

### Histology

Tissue was collected at the following developmental time points: post-natal day (P) 7, 14, 16, 28-30. Mice were anaesthetized by I.P. injection of 100 mg/kg Ketamine /20 mg/kg Xylazine mix (obtained from Comparative Medicine program at TAMU) and transcardially perfused with PBS, then 4% PFA at room temp. Brains were removed and incubated in 4% PFA overnight at 4C, then washed 3 X 5 min with PBS, and cryoprotected in 30% sucrose for 2-3 days, before being embedded in TFM media (VWR # 100496-345), frozen in dry ice-ethanol slurry solution, and stored at -80C until use. Brains were sectioned using a cryostat (Leica CM1950) in sagittal or coronal orientations depending on experimental needs at a slice thickness of 18-20 µm, or 100 µm for morphological studies (Fig. 6). Sections were either mounted on Superfrost plus slides (Fisher #1255015) or kept in PBS (floating sections) and either used immediately for histological procedures (Immunohistochemistry) or stored at -80C /4C for later use. 3-5 mice from each sex and age group were used. For each mouse, 2-3 sections were imaged and analyzed.

### Western Blot

Mice were anaesthetized by I.P. injection of 100 mg/kg Ketamine /20 mg/kg Xylazine mix (obtained from Comparative Medicine program at TAMU) and then decapitated. Brains were rapidly removed and the VC dissected in ice-cold PBS, flash frozen and kept at – 80°C until use. 2-4 samples for each condition were analyzed.

## IMMUNOHISTOCHEMISTRY

Slide mounted or free-floating sections were blocked for 1 hour at room temperature in blocking buffer containing antibody buffer (100 mM L-lysine and 0.3% Triton X-100 in PBS) supplemented with 10% heat-inactivated normal goat serum. Primary antibodies diluted in antibody buffer with 5% goat serum were incubated shaking overnight at 4°C. The following day, slide mounted sections were washed 3 X 5 min and free-floating sections were washed 3 X 15 min with PBS with 0.2% Triton X-100 and secondary antibodies conjugated to Alexa Fluor were applied for 2 hours at room temperature. Slides were mounted in SlowFade Gold media with DAPI (LifeTech #S36939), covered with 12 mm glass coverslip (Carolina Biological Supply Company # 633029) and sealed with clear nail polish. The following primary antibodies were used: Rb anti IP3R2 (Alomone labs #ACC-116, 1:250), Gp anti-VGLUT1 (Millipore #AB5905, 1:1000), Gp anti-VGLUT2 (Millipore #AB2251 1:1000), Rb anti-PSD95 (Fisher #516900 1:250), Gp anti-VGAT (Synaptic Systems #131004 1:250), Rb anti-Gephyrin (Synaptic Systems #147008 1:500), Rb anti-Nf200 (Millipore Sigma #N4142 1:400), Chk anti-GFP (Invitrogen A10262 1:1000), Rb anti-S100β (Abcam #AB52642, 1:100). The following secondary antibodies were used: Gt anti-Rb Alexa-488 (Invitrogen #A11043), Gt anti-Gp Alexa-555 (Invitrogen #A21435), Gt anti-Chk Alexa-488 (Invitrogen #A32931), Gt anti-Rb Alexa-594 (Invitrogen #A11037), Gt anti-Gp Alexa-594 (Invitrogen #A11076), Gt anti-Ms Alexa-647 (Invitrogen #A21236). All secondary antibodies were used at 1:500 dilution.

## MICROSCOPY AND IMAGING

### Fluorescent microscopy

Was performed to image the expression of IP3R2 (Fig. 1, Fig. S1), developmental changes in Nf200 (Fig. 2), and c-FOS expression (Fig. 4, Fig. S4) using a Leica THUNDER Imager with LED3 light source, and sCMOS camera (Leica DFC9000) at 20X magnification. Single plane images (2048 X 2048 pixels) containing the visual cortex were taken for IP3R2 validation and Nf200 studies (Fig.1, Fig. S1, Fig. 3). For c-FOS imaging experiments (Fig. 4, Fig. S4), 5 μm, 6 section z stacks (2048 X 2048 pixels) containing the visual cortex, hippocampus, dorsal lateral geniculate nucleus, or superior colliculus were acquired for each brain section separately. Example images are shown as a single z plane image from the same location in the stack for each genotype. Thunder processing (Leica LASX software) was performed using default parameters for single plane imaging (instant computational clearing) to increase resolution and image clarity in the same way for all images.

**Figure 1.**
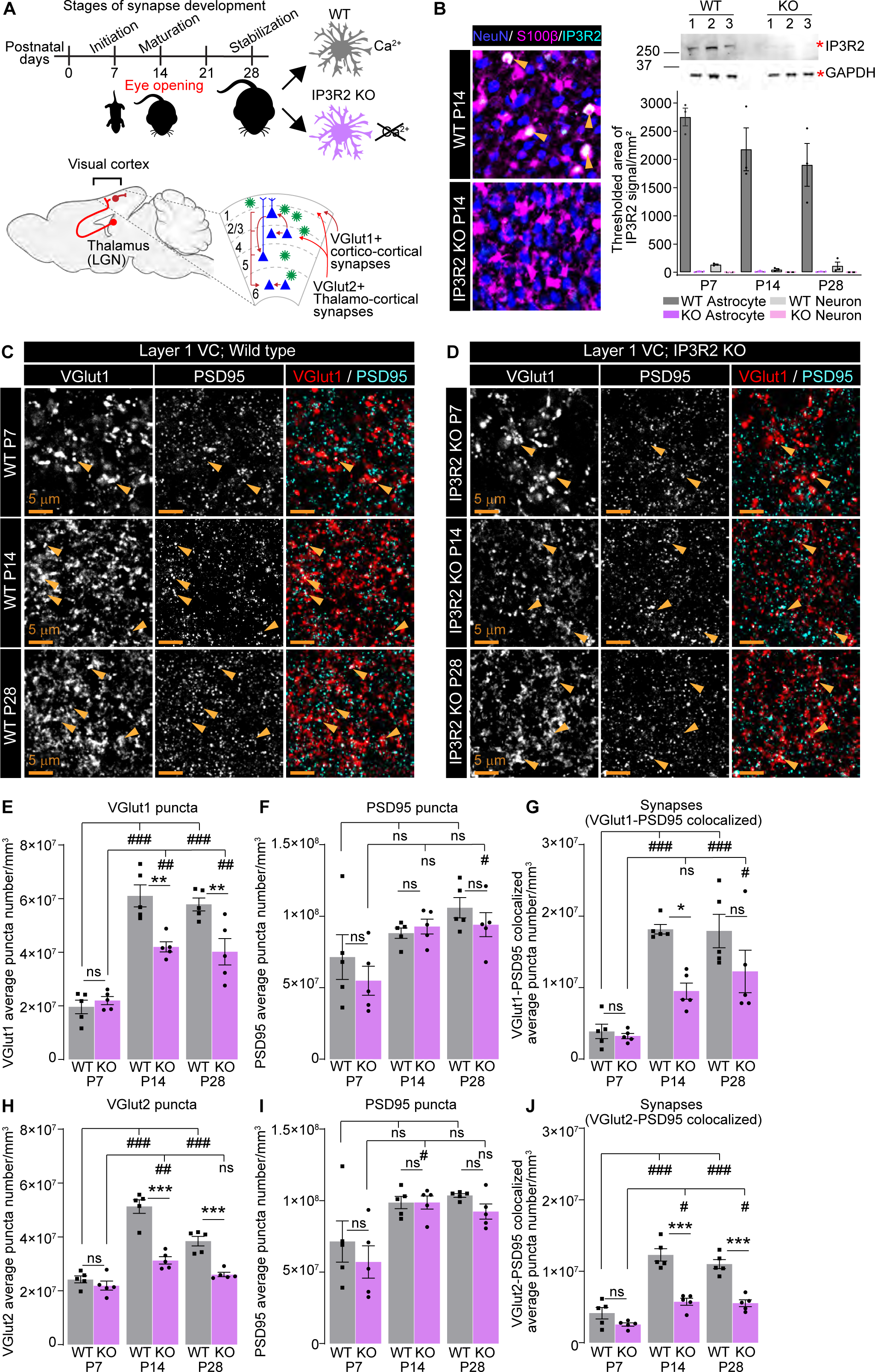
Glutamatergic synapse development is perturbed in IP3R2 KO mice VC. See also Fig S1. **A.** Schematic of experiment: Brain tissue is collected from WT and KO mice at P7, P14 and P28 corresponding to stages of synapse development; IHC to quantify synapses as indicated is performed in Layer 1 of the VC for VGLUT1-containing cortico-cortical synapses, and VGLUT2-containing thalamo-cortical synapses. **B.** Validation of IP3R2 KO by IHC and WB (top right panel). Example images of IP3R2 (cyan), astrocyte marker S100ꞵ (magenta) and neuronal marker Neun (blue) in the VC at P14 as labeled. Graph on the right is quantification of colocalized signal with each cell marker. IP3R2 signal is highly colocalized with astrocytes and not with neurons and is downregulated in KO VC. WB shows IP3R2 band (∼250KDa) and GAPDH (loading control, ∼36 KDa) in WT and KO at P14 as labeled. Numbers indicate samples from individual animals. See also Fig. S1A-D. **C-G.** Cortico-cortical VGLUT1-containing, synapses are reduced in IP3R2 KO VC at P14 and P28 but not P7. Example images of the presynaptic VGLUT1, postsynaptic PSD95 and merged (synapses) in each age and genotype as labeled (**C-D**) and quantification of individual synaptic proteins and synapse number per mm^3^ represented as colocalization between VGLUT1 and PSD95 for both genotypes (**E-F**) are shown. Single channel grayscale images on the left, merged images on the right. The number of VGLUT1 puncta and VGLUT1-containing synapses is reduced in KO (**E, G**), while PSD95 numbers are unaltered (**F**). **H-J.** Thalamo-cortical VGLUT2-containing synapses are reduced in IP3R2 KO VC at P14 and P28 but not P7. Quantification of individual synaptic proteins and synapse number per mm^3^ represented as colocalization between VGLUT2 and PSD95 for both genotypes are shown (see also Fig. S1E-F). The number of VGLUT2 puncta and VGLUT2-containing synapses is reduced in KO (**H, J**), while PSD95 numbers are unaltered (**I**). Plots show mean ± s.e.m. Squares and circles above each bar are average of signal in each mouse. Number of mice/ group (N) N=5. Scale bar = 5 μm. Arrowheads mark representative colocalized puncta. #,*P<0.05, ##, **P<0.01, ###, ***P<0.001. # indicates comparing age groups within each genotype by one-way ANOVA. Within each age, WT and KO comparison by t-test indicated by *. ns denotes Non-significant results (P>0.05).

**Figure 2.**
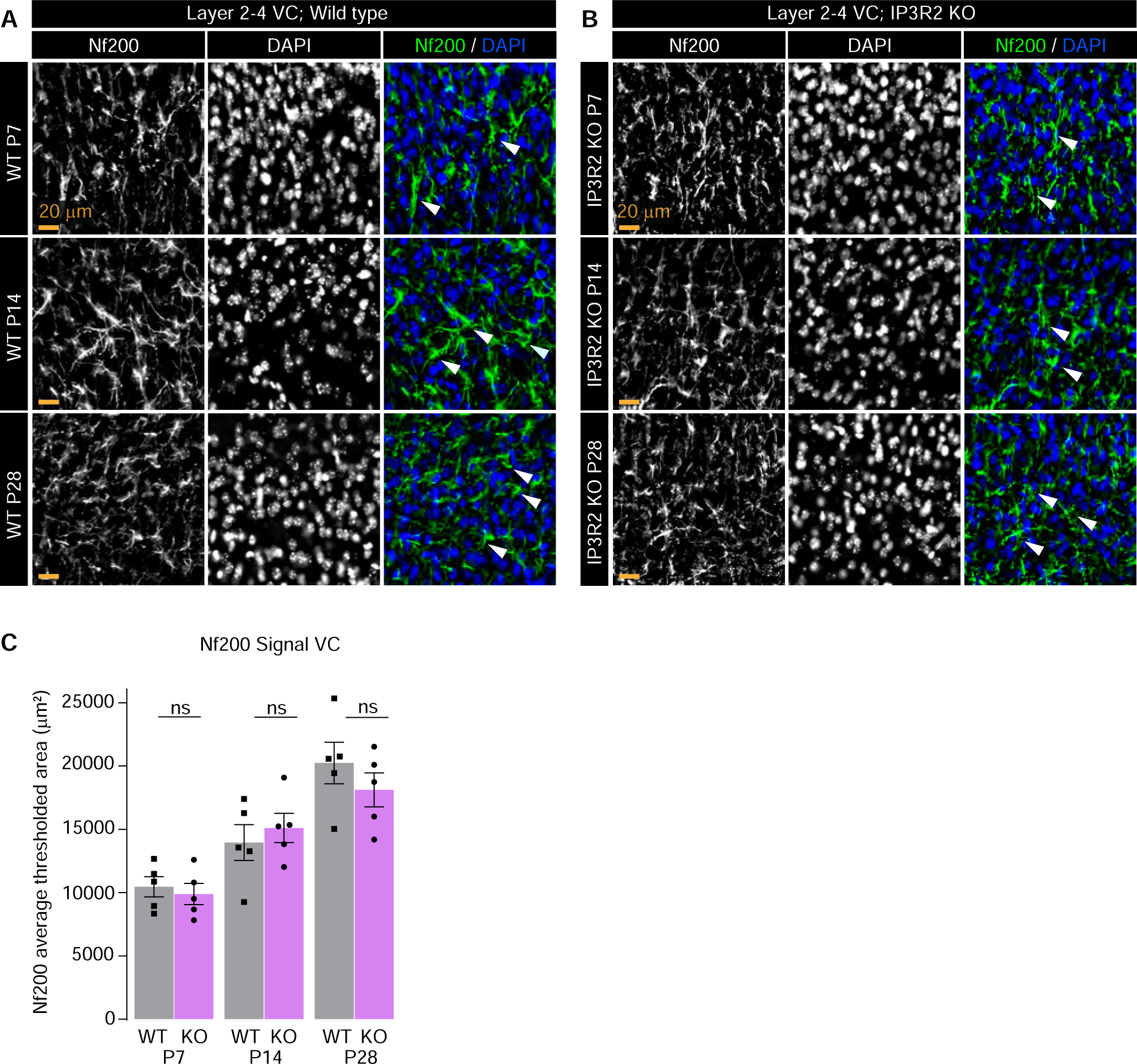
Axonal density is unperturbed in IP3R2 KO mice VC. A-B. Example images of the axonal marker Nf200 (green) and nuclear marker DAPI (blue) in each age and genotype as labeled. Single channel grayscale images on the left, merged images on the right. **C.** Quantification of Nf200 signal represented as thresholded area of signal for each age and both genotypes. No difference between WT and KO is observed at any age. Graph shows mean ± s.e.m. Squares and circles above each bar are average of signal in each mouse. Number of mice/ group (N) N=5. Scale bar = 20 μm. Arrowheads mark representative axons. ns denotes Non-significant (P>0.05) results comparing WT and KO signal within each age.

**Figure 3.**
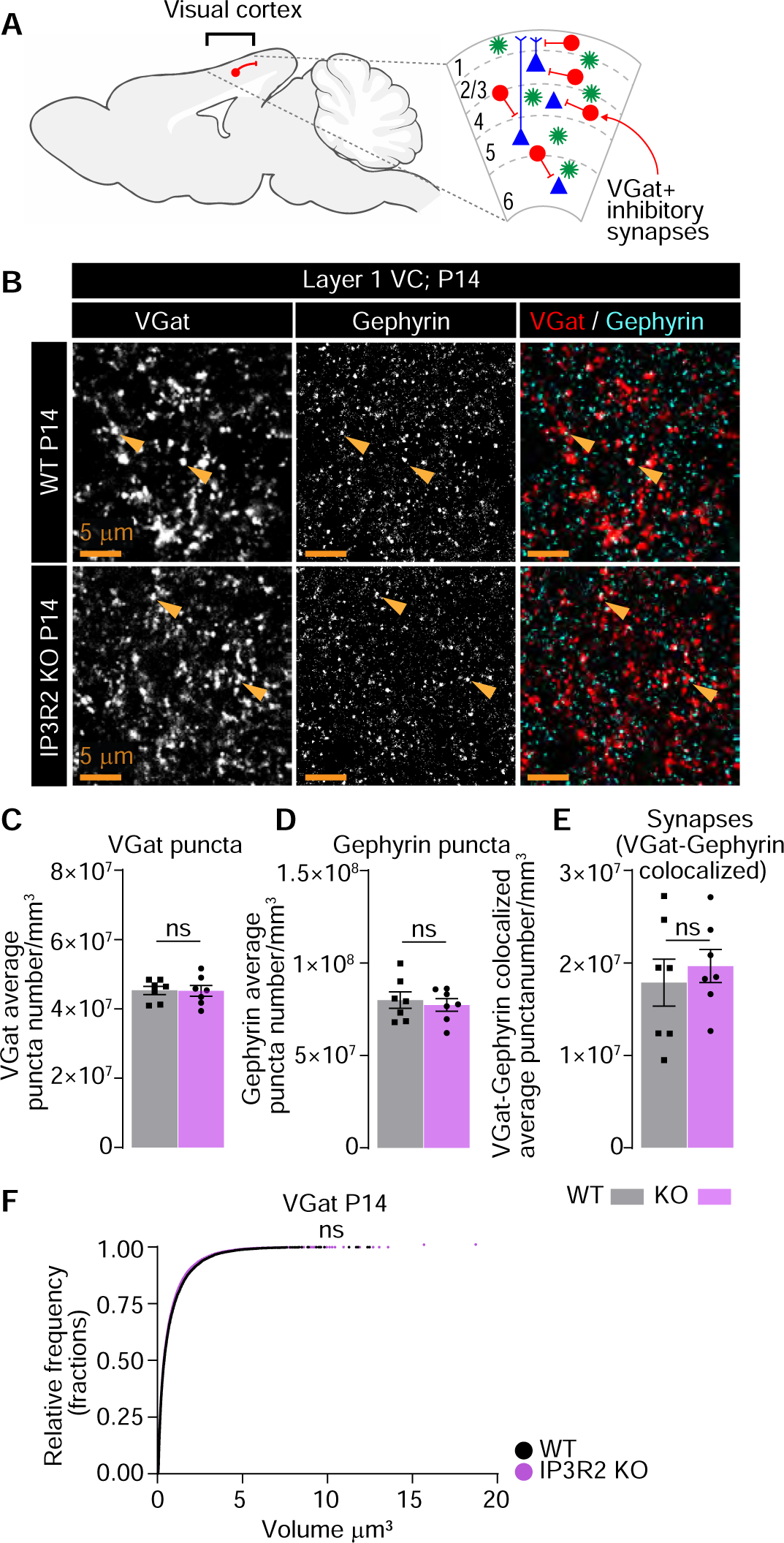
GABAergic synapses are unperturbed in IP3R2 KO mice VC at P14. **A.** Diagram depicting inhibitory neurons within the VC analyzed. **B-F.** GABAergic synapse numbers are not altered in IP3R2 KO VC at P14. Example images of the presynaptic VGAT, postsynaptic Gephyrin and merged (synapses) in each genotype as labeled (**B**) and quantification of individual synaptic proteins and synapse number per mm^3^ represented as colocalization between VGAT and Gephyrin for both genotypes (**C-E**) are shown. Single channel grayscale images on the left, merged images on the right. No difference is observed in any of the parameters compared. **F.** Cumulative distributions of volumes from 3D rendered images for VGAT per genotype as labeled. Bar graphs show mean ± s.e.m. Squares and circles above each bar are average of signal in each mouse. Number of mice/ group (N) N=5. Scale bar = 5 μm. Arrowheads mark representative colocalized puncta. ns denotes non-significant results (P>0.05) by t-test in C-E, and Kolmogorov-Smirnov test in F, comparing WT and KO groups.

**Figure 4.**
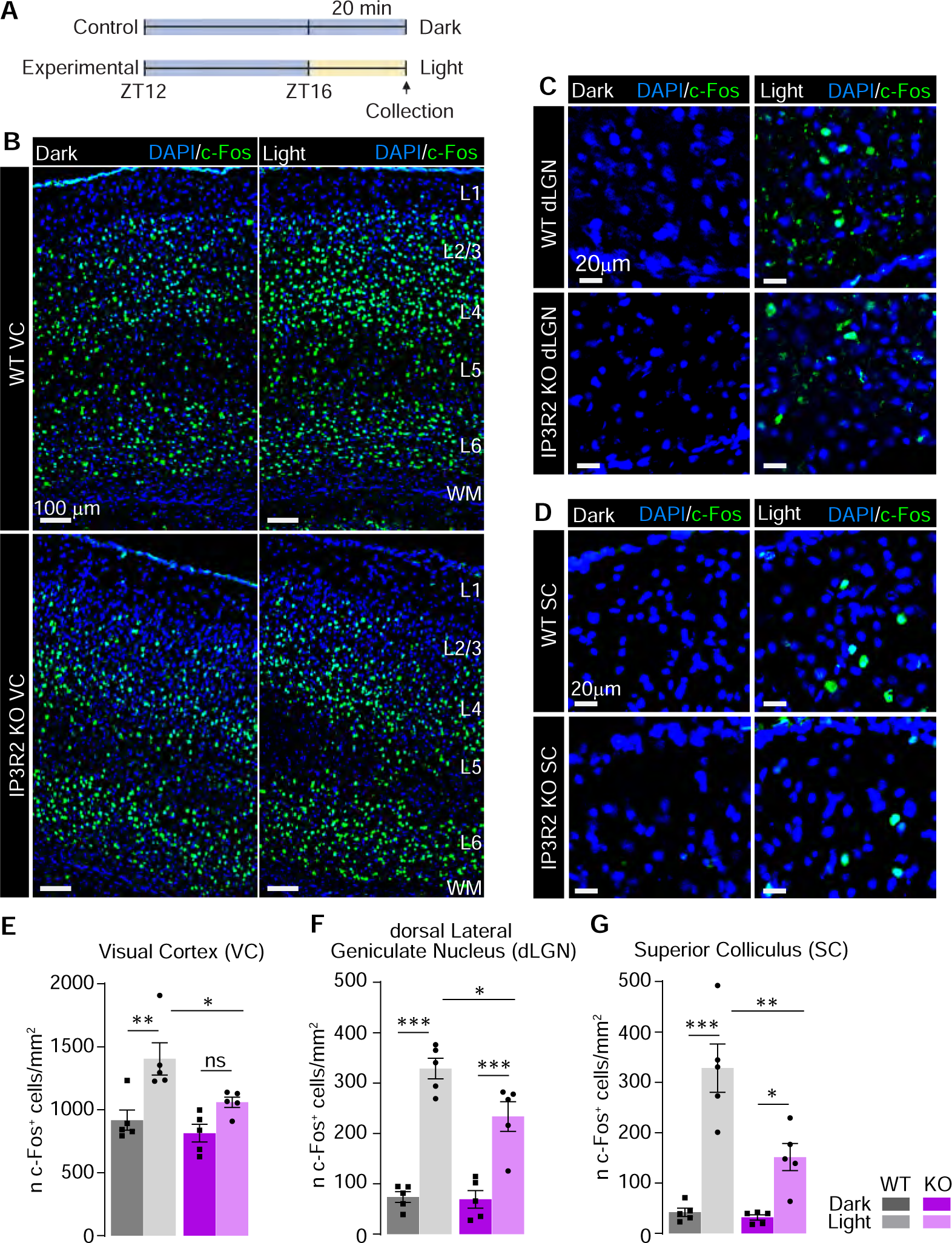
Light-evoked c-FOS expression is blunted in IP3R2 KO mice visual circuit. See also Fig. S4. **A.** Schematic of experimental paradigm. Following 4 hours of dark exposure (ZT12 marks lights off), mice were exposed to 20-minute light pulse, tissue collected immediately after for IHC analysis of c-FOS expression. **B, E.** Light evoked c-FOS levels are diminished in IP3R2 KO mice VC. Example images of c-FOS (green) and nuclear marker DAPI (blue) in each genotype as labeled in the dark or light exposed groups. Neuronal cortical layers are labeled on the right **E.** Quantification of B represented as c-FOS positive cell numbers per area. Light exposure produced a strong increase in the number of c-FOS positive cells in the WT, but not in the KO. **C-G.** Same as B, E, but for the dorsal lateral geniculate nucleus of the thalamus (dLGN; C, F) and the superior colliculus (SC; D, G). In both regions, light exposure produced a strong increase in the number of c-FOS positive cells in the WT, but to a lesser extent in the KO. Graphs show mean ± s.e.m. Squares and circles above each bar are average of signal in each mouse. Number of mice/ group (N) N=5. Scale bar in **B** = 100 μm; in **C-D** = 20 μm. *P<0.05, **P<0.01, ***P<0.001 by one-way ANOVA. ns denotes non-significant results (P>0.05).

### Confocal microscopy

was used to image synaptic proteins (Fig. 1, Fig. S1, Fig. S2, Fig. 3) and Lck-eGFP labeled astrocytes (Fig. 6, Fig. S6) Imaging was performed using a Zeiss LSM 900 upright confocal laser scanning microscope with Airyscan2. Synaptic imaging of VGLUT/PSD95 and VGAT/Gephyrin puncta (Fig.1, Fig. S1, Fig. 3) were acquired at 63X magnification. For each section, 1024 X 1024 pixels (101.4 X 101.4 X 2.79 µm) thick z stack image was obtained (pixel size 0.09 X 0.09 X 0.31 µm; 10 slices per 2.79 µm stack). Astrocyte morphological imaging (Fig. 6) was acquired at 63X magnification using the Airyscan2 module, 1834 X 1834 pixels (78.01 X 78.01 XY µm), and 27-47 µm stack to ensure imaging included the entire cell, pixel size 0.043 X 0.043 X 0.15 µm). Astrocytes located in layer 1 of the VC were selected for imaging if they did not directly connect to the pia (limitans) or if soma was present in layer 2. For validation of Lck-eGFP localization with astrocyte marker S100ꞵ (Fig. S6), 20X magnification was used to acquire 512 X 512 pixels (247.35 X 247.35 X15 µm) z stacks (pixel size 0.48 X 0.48 X 0.31 µm). Example images of synaptic proteins (Fig. 1, Fig. S1, Fig. 3) are shown as a single z plane image from the same location for each stack in the genotype, while example images for Lck-eGFP labeled astrocytes are shown as maximal intensity projections.

## IMAGE ANALYSIS AND QUANTIFICATION

Image analysis was done with ImageJ (NIH) or Imaris (Bitplane) software as described: *Quantification of pre-/postsynaptic puncta and synapses*: (Fig. 1, Fig. S1, Fig. S2, Fig. 3) was performed as previously described^4^ on 3D images using Imaris software (Bitplane). The surface function was used to create volumes for VGLUT or VGAT signal, while the spots function was used to render PSD95 or Gephyrin. Total number of surfaces/spots was obtained to quantify pre- and post-synaptic puncta count, and synapses were quantified by counting post-synaptic ‘spots’ located ≤0.25 µm of a presynaptic surface. Number of total and colocalized puncta were obtained and compared between the experimental groups. A minimum of three sections per mouse were imaged for each brain region, and the experiment was repeated in at least five WT and KO pairs. Volumes were computed from the surfaces generated for VGLUT1, VGLUT2 or VGAT and exported from Imaris.

### Quantification of IP3R2 signal

(Fig. 1, Fig. S1) Images labeled with IP3R2 and S100β (to label astrocytes) or Neun (to label neurons) were analyzed using a semi-automated custom-made macro in ImageJ. For each image, VCs were manually cropped and saved to a new file. For each of the different signals (IP3R2, S100β or Neun), images were thresholded in the same way for each section, and the IP3R2 signal area within S100β or Neun ROIs was calculated as thresholded area. Resulting thresholded areas of the IP3R2 signal were normalized to the total area to obtain thresholded area per mm^2^.

### Quantification of axonal density

(Fig. 2) Images labeled with Nf200 (to label axons) and DAPI (to label nuclei) were analyzed using a semi-automated custom-made macro in ImageJ. For each image, VCs were manually cropped and saved to a new file. Images were thresholded in the same way for each section for Nf200 signal, and the ‘analyze particles’ function was used with a size range of 10-150 pixels to quantify DAPI. Resulting average thresholded area was normalized to the total area analyzed to obtain thresholded area per μm^2^, and cell counts were also normalized to the total area to obtain the number of cells per mm^2^.

### Quantification of c-FOS

(Fig. 4) Images labeled with anti-c-Fos antibody and DAPI (to label nuclei) were analyzed using a semi-automated custom-made macro in ImageJ. Maximum projection images were generated from z stacks, cropped by cortical layer (VC) or region (dLGN, SC, CA1, DG) and saved as individual files for analysis. Sections containing dLGN were co-stained for VGLUT2 to identify regional boundaries. Each channel was thresholded using the dark-background auto-threshold method, followed by manual adjustment by the user. For c-FOS, particles within a size range of 25–350 pixels were detected, and the total count was measured within ROIs. For DAPI, particles sized 10–150 pixels were segmented following watershed separation, and nuclei were counted.

### Quantification of astrocyte morphology

(Fig. 6) was performed on 3D images using Imaris. The surface function was used to create volumes via absolute intensity and cropped to remove partial signal from neighboring astrocytes within the field of view. Remaining surfaces were unified to generate outputs for total volume and area as well as sphericity and ellipticity. A minimum of 5 cells from 3 different sections in at least 5 WT and KO pairs were imaged and analyzed.

### Quantification of GfaABC1D-Lck-GFP colocalization with S100β

(Fig. S6) Images containing astrocytes expressing Lck-eGFP were co-labeled with antibody for S100β (to mark astrocytes) and analyzed using a semi-automated custom-made macro in ImageJ. For each of the different signals (Lck-eGFP, S100β), images were thresholded in the same way for each section, and the “cell counter” tool was used for manual annotation of cell types. Colocalized cells were counted using a separate counter type. Resulting cell and colocalization counts were analyzed to obtain % overlapping cells.

## INTRACEREBROVENTRICULAR (ICV) AAV INJECTION

Was performed as previously described^34–36^. P1 mouse pups were removed from their home cage and placed in a warmed holding cage. Pups were individually cryo-anaesthetized in ice for 5 minutes (placed inside a glove) before being transferred to a surgical pad over a back-lit surface. The ventricles were identified by measuring the distance 2/5^ths^ from the lambda suture to the eye. 1-2 µL of AAV5-GfaABC1D-Lck-GFP (VectorBuilder, #VB240226-1200bvh) diluted to 2 X 10^9^ gc/mL in saline was drawn into the barrel of a 10 µl glass syringe (Hamilton #7653-01) fitted with a 32G injection needle (Hamilton #7803-04). With the pup’s head upright and secured by hand, the injection site was swabbed with 70% ethanol and the tip of the needle was positioned perpendicular to the injection site and inserted 3 mm before the plunger was manually depressed. The syringe was held in place for 10 seconds to avoid ejection, and then carefully withdrawn. This procedure was performed for both ventricles. After injection, pups were returned to the warmed holding cage for immediate recovery, then to their home cage.

## LIGHT PULSE ASSAY

P16 mice (aged to ensure eyes were open for 24 hours before light pulse experiments) were habituated in cages placed within darkened boxes (32W X 38L X 32 X 54.6H cm) for 30 minutes prior to experimentation, then mice were moved (individually) to either a control cage within a darkened box, or an experimental cage in a separate box fitted with 3 white LED light strips (110 lumen each). Experimental mice were subjected to 330 lumen (∼2713 lux) light pulse for 20 minutes, followed by immediate anesthesia by Ketamine/Xylazine injection and perfusion as described (see Mouse tissue section). Control mice remained in dark condition for 20 minutes before tissue collection. All mice were collected between ZT16 – ZT17 after four hours of sustained darkness (ZT12-ZT16) under standard housing conditions.

## VISUALLY EVOKED DEFENSIVE BEHAVIORAL ASSAY

### Behavioral testing setup and testing schedule

The test was performed according to published protocols with modifications^37,38^. The setup consists of a rectangular arena (17W X 40L X 20H cm) with a plastic hut (11 cm in diameter, 5 cm height) placed in one corner of the cage serving as shelter for mice to escape to. The hut used is identical to that used in the mouse home cages to facilitate quicker familiarization and habituation to the testing environment. A computer monitor is mounted 40 cm above the arena floor, connected to a computer through which the looming stimulus is delivered. A video camera is placed above the cage to record mouse movement during the stimulus for offline analysis. The test was administered over 3 consecutive days and consisted of a habituation day (day 0), followed by 2 consecutive testing days (see Fig. 5A). During habituation, mice are placed in the testing arena one at a time, and allowed to explore for 13 minutes, without the looming stimulus played. On each Testing day, mice are placed in the arena and subjected to 5 trials of the looming stimulus. The first stimulus is played after a minimum of 3 minutes of free exploration, with a minimum of 1 minute interval between subsequent stimuli, during which time mice resume exploratory behavior. Mice spend a total of 13 minutes in the arena on testing days, then are returned to home cages even if they did not receive 5 trials. Stimulus is played only if a mouse is outside the shelter and engaged in exploratory behavior. Test day 2 is identical to Test day 1, occurring ∼24 hours later. Mice are returned to the vivarium between testing days.

**Figure 5.**
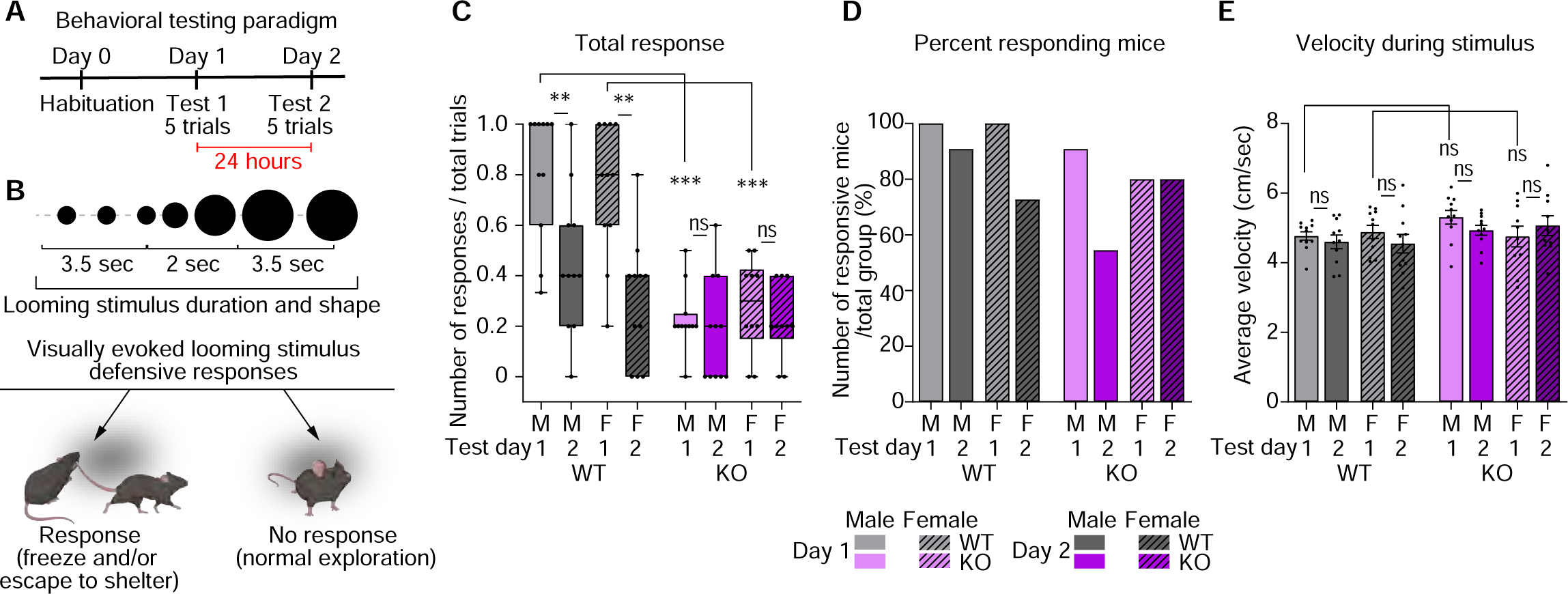
Visually evoked defensive behavior is disrupted in IP3R2 KO mice. See also Fig. S5. **A.** Diagram of the behavioral experiment including habituation and 2 testing days performed 24 hours apart. Each testing day includes 5 trials of the looming stimulus. Mice are naïve to the stimulus on Test day 1 and retested on Test day 2 (experienced). **B.** Schematic of the looming stimulus shape and duration (top) and the types of defensive responses analyzed (bottom). **C.** Average defensive responses (of any type) across genotypes, sexes, and experimental days. WT mice of both sexes exhibit robust response to the looming stimulus on Test Day1 and reduced response when retested (Test day 2). KO mice responses are strongly downregulated showing no difference between testing days. **D.** Number of responders represented as percentage of mice responding to at least one trial per day out of total group number. All groups had similar percentage of mice responding to the stimulus. **E.** Velocity averaged across the entire 9 seconds of the stimulus shows similar kinematic parameters for all sexes and genotypes. In **C**, graph shows box with range, line is median. Circles on each box are responses of individual mice. In **D**, graph shows percentage of responding mice out of total group. In **E**, graph shows mean ± s.e.m. Squares and circles above each bar are average of velocity for each mouse. Number of mice (N): 8-10/ per sex/ genotype. *P ≤ 0.05, **P<0.01, ***P<0.001 by Mann-Whitney test. ns denotes non-significant results (P>0.05).

### The looming stimulus

A 9 second video showing an expanding black disc (2-20cm in diameter) on a white background is played to mimic a looming threat (file available from^37^). The stimulus begins when a black disc (2 cm) appears on the computer screen (∼ 3.5 seconds), rapidly expands to 20 cm diameter (∼2 seconds) and remains fully expanded for ∼3.5 seconds. This stimulus has been thoroughly characterized in previous studies to elicit robust defensive responses in mice^37,38^.

### Quantification of responses and kinematics

1. <u>Defensive behavioral scoring</u> – was done by an experimenter either live during the testing, or offline from recorded videos. For each stimulus, 4 types of responses are recorded: “Freeze” – a mouse’s snout and body is immobile for a minimum of 1 second; “Escape” – the mouse is moving towards and entering the shelter; “Freeze+Escape” – a period of immobility followed by movement and entering the shelter; “No response” – mouse continues exploratory behavior or movement with no directed change due to stimulus.
2. <u>Kinematics analysis</u> – was done by using DeepLabCut^39,40^ trained custom pose estimation model for tracking mice during behavioral experiments, which was then applied to the video recordings to generate framewise X-Y coordinates for each frame. The resulting data was used as input by BehaviorDEPOT^41^ to calculate the velocities (computed using the Pythagorean Theorem: √[(dx/dt)² + (dy/dt)²]). Custom MATLAB and Python scripts were used to extract kinematic data from the entire recording and from specific frame ranges corresponding to the presentation of the looming stimulus, respectively. The scripts (<u>GitHub_scripts</u>) and the DeepLabCut model (<u>DLC model</u>) can be accessed through the provided links.

### Exclusion criteria

Mice which missed more than 2 stimuli (e.g. remaining inside the shelter/failure to explore) on either testing day were removed from the study.

## WESTERN BLOT

Samples were heated in reducing loading dye (Thermo Scientific # 39000) for 45 min at 55°C. For tissue lysates, 5 µg/lane was loaded. Samples were resolved on Bolt 4–12% Bis-Tris gels (Invitrogen #NW04125BOX) for 1 hour and 15 min at 150 V. Proteins were transferred to Immobilon-FL PVDF membranes (Millipore #IPFL00010) at 20 V for 1 hr, then blocked in 1% casein (Bio-Rad #1610782) in TBS (Bioworld #10530027-2) blocking buffer for 1 hr at room temperature on a shaker. Primary antibodies were applied overnight at 4C diluted in blocking buffer. The antibodies used were Rb anti-IP3R2 (Alomone labs # ACC-116, 1:500), and Ms anti-GAPDH (Cell Signaling Technology # 97166T, 1:2000). The next day, membranes were washed 3 X 10 min with TBS-0.1% Tween (Promega # PRH5152) and secondary antibodies Gt anti-Rb Alexa-680 (Invitrogen # A21109) and Gt anti-Ms-Alexa-800 (Invitrogen # PIA32730) were applied for 2 hr at room temperature (dilution 1:10,000). Bands were visualized using the LI-COR Odyssey CLx Infrared Imager and band intensity analyzed using the LI-COR Image Studio software.

## STATISTICAL ANALYSIS

Data is shown either as mean ± s.e.m, median and range, or percentages, as indicated in each Figure legend. Statistical analysis was performed using Prism software (Graphpad). Multiple group comparisons were done using one-way analysis of variance (ANOVA) with post hoc Tukey’s or Dunn’s tests, while pairwise comparisons were done by t-test for normally distributed data or Kruskal-Wallis ANOVA on ranks for multiple comparison with post hoc Dunn’s test, and Mann-Whitney rank sum test for pairwise comparison for data that did not exhibit normal distribution. Each dataset was tested for normality (with Shapiro-Wilks test) to ensure correct statistical test is used. Fisher’s exact test was used to compare percentages of responding mice (Fig. 5, Fig. S5). Two sample Kolmogorov-Smirnov test was used to assess differences between cumulative frequency distributions. Analysis was completed blind to genotype. Sample sizes, statistical tests used, and significance are presented in each Figure and Figure legend. P Value ≤ 0.05 was considered statistically significant.

## RESULTS

### Vesicular glutamate transporter and synapse numbers are developmentally reduced in the visual cortex of IP3R2 knockout mice

Astrocytic intracellular Ca^2+^ fluctuations are fundamental to their regulatory roles at the neuronal synapse. While these Ca^2+^signals are characterized by a diversity of forms and origins^42,43^, store-released Ca^2+^mediated by the astrocytic IP3R2 couples synaptic neurotransmitter release to changes in synapse regulating genes in astrocytes^4,44^. We previously identified that mice lacking IP3R2, which is essential for astrocytic store-released Ca^2+^signaling, have disrupted expression of synapse regulating genes and decreased numbers of vesicular glutamate transporters at postnatal day (P) 14^4^. To investigate whether synapses in IP3R2 knockout (KO) mice are broadly dysregulated across development, we profiled the expression of pre- and post-synaptic proteins in layer 1 of the visual cortex (VC) of wild-type (WT) and IP3R2 KO mice at P7, P14, and P28 which correspond to cortical synaptogenic onset, peak, and stabilization respectively^2^ (Fig. 1A). We first validated the deletion of IP3R2 at the protein level using immunohistochemistry (IHC) and western blot across the 3 developmental stages studied (P7, P14, P28) (Fig. 1B, Fig. S1A-D). For IHC, brain sections were co-labeled with astrocyte specific marker S100β and neuronal specific marker Neun to determine cell-type specificity of IP3R2 expression. Our results show colocalization of IP3R2 signal with S100β but not with Neun, confirming the astrocytic source of this receptor within the developing VC gray matter (shown as mean ± s.e.m. [thresholded area µm^2^]: S100β – [P7] WT: 2752 ±156.7, KO: 11.05 ±5.518; [P14] WT: 2180 ±380.4, KO:13.48 ±7.38; [P28] WT: 1905 ±379.9, KO: 10.71 ±3.198; Neun – [P7] WT: 135.4 ±10.31, KO: 1.255 ±1.205; [P14] WT: 47.31 ±15.91, KO: 3.66 ±1.084; [P28] WT: 112.6 ±68.56, KO: 0.3501 ±0.1442; Fig. 1B). IP3R2 protein levels are strongly reduced in tissue from IP3R2 KO (by both IHC and WB) mice at all ages tested confirming this model’s usage for our experiments (WB data is shown as mean + s.e.m normalized to GAPDH: [P7] WT: 0.1359 ±0.0072, KO: 0.0025 ±0.0012; [P14] WT: 0.1534 ±0.0186, KO: 0.02741 ±0.0095; [P28] WT: 0.0620 ±0.0033, KO: 0.0101 ±0.0005; Fig. 1B, Fig. S1D).

To analyze how glutamatergic synapse development is impacted by IP3R2 KO, we used IHC to quantify synapses of 2 glutamatergic circuits within the VC, the presynaptic vesicular glutamate transporters (VGLUT1) specific cortico-cortical circuit and vesicular glutamate transporters (VGLUT2) specific thalamo-cortical circuit together with the excitatory postsynaptic scaffolding protein postsynaptic density protein 95 (PSD95). We quantified both the numbers of pre- and postsynaptic puncta and their colocalizations to calculate synapse number across these developmental stages^4^ (Fig. 1C-J, Fig. S1E-F).

We observed a significant decrease in VGLUT1 (∼30%) and VGLUT2 (∼40%) at P14 which persisted to P28 in VC sections from IP3R2 KO mice (∼30% for both proteins) (data shown as mean ± s.e.m. [puncta number per mm³]: P14: VGLUT1 – [P14] WT: 6.10E7 ±4.08E6, KO: 4.20E7 ±1.89E6, [P28] WT: 5.78E7 ±2.40E6, KO: 4.02E7 ±4.92E6; VGLUT2 – [P14] WT: 4.92E7 ±2.43E6, KO: 2.99E7 ±1.32E6, [P28] WT: 3.68E7 ±1.72E6, KO: 2.49E7 ±7.76E5; Fig. 1E, H). Synapse numbers were also decreased for both VGLUT1 and VGLUT2 containing projections at P14 (∼41% and 35% respectively) and P28 (∼32% and 50% respectively) (VGLUT1-PSD95 – [P14] WT: 1.82E7 ±6.72E5, KO: 9.51E6 ±1.13E6; [P28] WT: 1.79E7 ±2.34E6, KO: 1.23E7 ±2.98E6; VGLUT2-PSD95 – [P14] WT: 1.23E7 ±8.69E5, KO: 5.73E6 ±4.93E5; [P28] WT: 1.10E7 ±6.39E5, KO: 5.53E6 ±4.77E5; Fig.1G, J). On the other hand, no difference in VGLUT numbers or synapses were observed at P7 (VGLUT1 – WT: 1.96E7 ±2.53E6, KO: 2.20E7 ±1.49E6; VGLUT2 – WT: 2.31E7 ±1.25E6, KO: 2.09E7 ±1.63E6; Fig. 1E, H. Synapses (colocalized): VGLUT1-PSD95 – WT: 3.87E6 ±1.01E6, KO: 3.22E6 ±3.71E5; VGLUT2-PSD95 – WT: 4.14E6 ±7.92E5, KO: 2.55E6 ±2.41E5; Fig. 1G, J). The observed synaptic deficits were driven by the reduction in presynaptic proteins, as levels of PSD95 were unchanged between WT and IP3R2 KO mice at all ages tested ([P7] WT: 7.14E7 ±1.57E7; 7.14E7 ±1.44E7, KO: 5.49E7 ±1.01E7; 5.71E7 ±1.13E7; Fig. 1F. [P14] WT: 8.81E7 ±3.64E6; 9.86E7 ±4.29E6, KO: 9.28E7 ±5.20E6; 9.87E7 ±4.59E6, [P28] WT: 1.06E8 ±7.09E6; 1.04E8 ±1.25E6, KO: 9.41E7 ±8.42E6; 9.24E7 ±5.30E6; Fig. 1F, I).

To further analyze how VGLUTs are disrupted in IP3R2 KO mice we calculated the volumes of individual puncta (from 3D rendering, see Methods) and computed the cumulative frequency of VGLUT puncta volumes for WT and IP3R2 KO mice each age (Fig. S2A-F). We observed that while at P7, WT and IP3R2 KO mice had similar VGLUT1 and VGLUT2 volumes (data shown as mean ± s.e.m. [volume µm³]: VGLUT1 – WT: 0.8971 ±0.01215, KO: 0.8491 ±0.01103; VGLUT2 – WT: 0.8392 ±0.01086, KO: 0.7844 ±0.01119; Fig. S2A, D), P14 and P28 VGLUT1 and VGLUT2 volumes were significantly reduced in IP3R2 KO mice relative to WT mice, by ∼40%, and ∼35% respectively at P14, and ∼14% and ∼20% at P28. (VGLUT1 – [P14] WT: 1.120 ±0.0096, KO: 0.6654 ±0.0066; VGLUT2 – WT: 0.7910 ±0.0082, KO: 0.5154 ±0.0072. VGLUT1 – [P28] WT: 1.006 ±0.011, KO: 0.8637 ±0.012; VGLUT2 – WT: 0.8093 ±0.0092, KO: 0.6484 ±0.0088; Fig. S2B-C, E-F). These data suggest that IP3R2 mediated Ca^2+^ signaling is necessary for the developmental maintenance, but not initiation, of glutamatergic synapses in the VC.

### Axon density and cell number are unaffected by IP3R2 knockout

The observed decrease in VGLUT1 and VGLUT2 immunoreactivity could also be indicative of reduced cortical and thalamic innervation to the VC. To assess this possibility, we immunolabeled VC sections from WT and IP3R2 KO mice at P7, P14, and P28 for the axonal intermediate filament protein neurofilament 200 (Nf200) to quantify axon density^45^(Fig. 2). Axon density increased steadily across developmental stages supporting previous studies^46,47^, with no changes observed between WT and IP3R2 KO mice at any of the timepoints assessed (data shown as mean ± s.e.m. [thresholded area µm^2^]: [P7] WT: 10460 ±803.0, KO: 9883 ±836.3, [P14] WT: 13953 ±1414, KO: 15108 ±1163, [P28] WT: 20239 ±1646, KO: 18117 ±1339; Fig. 2C). We also tested the possibility that reductions in synapse number in IP3R2 KO mice could be due to a general decrease in cell survival or proliferation across development by quantifying the number of DAPI positive cells across the visual cortex at P7, P14, and P28 and, in accordance with our previously published data at P14^4^, observed no difference between WT and IP3R2 mice (data shown as mean ± s.e.m. [cell number per mm^2^]: [P7] WT: 4504 ±253.7, KO: 4186 ±120.5, [P14] WT: 2274 ±99.58, KO: 2262 ±101.2, [P28] WT: 2275 ±221.1, KO: 2221 ±230.1; Fig. S2G). These results suggest that the developmental synaptic protein deficiencies observed in IP3R2 KO mice are synapse-specific, and not due to a decrease in cell number or axonal innervation in the VC.

### Inhibitory synapse development in the visual cortex is unaltered by IP3R2 knockout

Mouse VC is composed of ∼80% glutamatergic and 20% GABAergic neurons^48^.To test whether the observed synaptic deficits are specific to excitatory circuits we used IHC to label pre-synaptic vesicular GABA transporters (VGAT) and the inhibitory post-synaptic scaffolding protein Gephyrin in WT and IP3R2 KO mice at P14 (Fig. 3A, B), the timepoint at which we observed the greatest deficits in glutamatergic synapses (Fig. 1C-J). We found no change in the number of VGAT puncta (shown as mean ± s.e.m. [puncta number per mm³]: WT: 4.53E7 ±1.18E6, KO: 4.52E7 ±1.58E6; Fig. 3C), Gephyrin puncta (WT: 8.00E7 ±4.40E6, KO: 7.73E7 ±3.36E6; Fig. 3D), or colocalization (WT: 1.79E7 ±2.54E6, KO: 1.20E7 ±1.79E6; Fig. 3E), and no change in VGAT volumes between the genotypes (shown as mean ± s.e.m. [volume µm³]: WT: 0.8051 ±0.0073, KO: 0.8059 ±0.0069; Fig. 3F). These results demonstrate that synaptic disruptions in IP3R2 KO mice are specific to glutamatergic terminals in the VC.

### Visually evoked neuronal immediate early gene c-FOS expression is blunted in IP3R2 knockout mice VC

Reduction in excitatory synaptic proteins in the visual cortex could contribute to deficits in visually evoked neuronal activity, and it has been shown that astrocyte Ca^2+^ fluctuation is important for the modulation of neuronal activity^4,42^. To test whether disruption of store released Ca^2+^ in astrocytes affects the activation of neurons in the VC, we examined activity dependent expression of the protein product of the immediate early gene *c-fos* (protein name c-FOS) following a brief light pulse stimulus (Fig. 4 A) in 16-day old mice (to ensure mice had at least 24 hours of processing visual information following eye opening at ∼P14). Fluorescent immunolabeling of c-FOS protein across the entire VC revealed that in WT mice, light stimulation induced a robust increase in the number of c-FOS positive cells (∼50%) compared to mice kept in the dark (shown as mean ± s.e.m. [cell number per mm^2^]: WT – dark: 917.4 ±80.38, light: 1404 ±127.9; Fig. 4B, E). When separately analyzing each of the cortical neuronal layers (L)^4^, this increase was strongest in L2-4, which receive visual input from the thalamus (L4) and integrate input response (L2-3) (WT – [L2/3] dark: 892.7 ±63.37, light: 1649 ±107.0, [L4] dark:1512 ±143.6, light: 2090 ±170.9; Fig. S4B, C). In the IP3R2 KO mice VC, this effect was blunted, with an increase of c-FOS positive cells of ∼30% (KO – dark: 814.2 ±69.86, light: 1059 ±41.08; Fig. 4B, E), and layer specific c-FOS induction decreases relative to WT mice in L2/3 (∼30% lower in KO mice) and L4 (∼20% lower in KO mice) (KO – [L2-3] dark: 764.1 ±112.1, light: 1135 ±98.36, [L4] dark:1424 ±77.12, light: 1676 ±79.47; Fig. S4B, C). While no significant difference was observed between the genotypes in L1, L5, and L6 (WT – [L1] dark: 41.81 ±5.92 light: 80.90 ± 27.21, KO – dark: 31.59 ±7.42, light: 82.92 ±16.74; [L5] WT – dark: 770.5 ±81.37, light: 1180 ±164.0, KO – dark: 657.8 ±92.57, light: 924.5 ±78.37; [L6] WT – dark: 1247 ±131.5, light: 1654 ±168.9, KO – dark: 1088 ±96.10, light: 1313 ±69.65; Fig. S4A, D, E). The observed effects were not due to changes in cell number as evident from similar DAPI-positive signal across the genotypes (data shown as mean ± s.e.m. [cell number per mm^2^]: [L1] WT – dark: 1360 ±42.84, light: 1430 ±83.73; KO – dark: 1420 ±70.53, light: 1281 ±63.49; [L2/3] WT – dark: 2879 ±113.5, light: 3069 ±212.2; KO – dark: 2911 ±86.31, light: 2790 ±201.8; [L4] WT – dark: 3377 ±324.0, light: 3801 ±283.6; KO – dark: 3162 ±190.8, light: 2916 ±252.3; [L5] WT – dark: 2523 ±202.8, light: 2201 ±182.4; KO – dark: 2678 ±220.4, light: 2414 ±348.7 [L6] WT – dark: 3724 ±281.0, light: 3617 ±225.4; KO – dark: 3444 ±343.9, light: 3207 ±279.1; Fig. S4I-M) To test whether the observed attenuated activity in the VC reflected upstream changes in subcortical visual processing, we examined c-FOS expression in the dorsal lateral geniculate nucleus (dLGN) and superior colliculus (SC) (Fig. 4 C, D) which receive direct retinal input and contribute to VC function^49–52^. WT mice exhibited dramatic increases in c-FOS positive cell numbers in both the dLGN (∼340%) and SC (∼680%) (WT – [LGN] dark: 73.99 ±10.67, light: 328.9 ±20.64, [SC] dark: 41.93 ±8.107, light: 328.2 ±48.14; Fig. 4 F, G). However, this increase was blunted in IP3R2 KO brains showing ∼320% increase in the dLGN and ∼380% in the SC (KO – [LGN] dark: 69.13 ±17.36, light: 233.8 ±29.52 [SC] dark: 31.80 ±5.310, light: 151.5 ±26.94; Fig. 4 F, G), in both regions, the numbers of light-induced c-FOS positive cells was significantly lower than in WT mice (Fig. 4F, G). As with the VC results, these c-FOS changes were not due to differences in cell numbers between WT and IP3R2 KO mice as evident from DAPI signal (WT – [LGN] dark: 3074 ±61.99, light: 3091 ±197.4; [SC] dark: 3427 ±196.8, light: 3463 ±196.1; KO – [LGN] dark: 2487 ±159.8, light: 2871 ±285.5; [SC] dark: 3240 ±134.6, light: 3400 ±146.6; Fig. S4P-Q). To ensure that the effects of the light pulse were vison-specific, we quantified the number of c-FOS positive cells in the CA1 and dentate gyrus (DG) regions of the hippocampus (Fig. S4H) and observed no significant changes in c-FOS protein induction following light pulse between the genotypes (WT – [CA1] dark: 1532 ±233.9, light: 1992 ±365.5 [DG] dark: 309.6 ±32.10, light: 369.4 ±37.41; KO – [CA1] 1324 ±254.0, light: 1642 ±145.0 [DG] dark: 295.0 ±44.34, light: 288.1 ±40.27; Fig. S4F-H). DAPI positive cell numbers were also unchanged between the genotypes (WT – [CA1] dark: 5812 ±439.8, light: 5194 ±473.5 [DG] dark: 4416 ±200.4, light: 4938 ±312.3; KO – [CA1] dark: 4683 ±527.3, light: 4418 ±510.3 [DG] dark: 4416 ±152.5, light: 4241 ±375.8; Fig. S4N, O). These results indicate that IP3R2 mediated astrocytic Ca^2+^ signaling plays an important role in facilitating visually evoked, but not basal, neuronal activity across the visual circuit brain regions.

### Visually evoked behavioral responses are reduced in IP3R2 knockout mice

Since we observed circuit level deficits in visually evoked neuronal activation, we next asked whether these findings translate to alterations in the development of behaviors which rely on these circuits for appropriate execution. Visually evoked defensive responses depend on an animal’s ability to perceive a threatening stimulus and integrate that information to coordinate an appropriate behavioral output^38,53^. We hypothesized that if IP3R2 KO mice experienced reduced or delayed activation of visual circuit neurons in response to salient stimuli at P16 (Fig. 4) and showed significantly reduced excitatory synapse number at P28 (Fig. 1G, J) when synaptogenesis is stabilized, they may exhibit disrupted behavioral responses to visual stimuli at later ages. To test this hypothesis, we subjected 4-week-old WT and IP3R2 KO mice (P30) to a visual looming threat and quantified their responses across two days, over which animals are considered naïve or adapted to the stimulus, respectively^37,54,55^ (Fig. 5A). Responses were scored (1 – response, 0 – no response) and categorized into four different classes: “Freeze” – mice exhibit immobility; “Escape” – mice move towards and enter the shelter; “Freeze+escape” – immobility followed by movement towards and entering the shelter; “No response” – mice continue exploratory behavior such as movement or sniffing (Fig. 5B).

WT mice of both sexes responded robustly to the looming stimulus on test day 1 (D1, naïve) and significantly reduced their responses on test day 2 (D2, experienced) (shown as median and [range]: WT – M D1: 1.0 [0.3-1.0], D2: 0.4 [0-1.0]; F D1: 0.8 [0.2-1.0], D2: 0.4 [0-0.8] Fig. 5C), demonstrating high sensitivity to the stimulus and rapid adaption upon re-exposure. In contrast, IP3R2 KO mice had highly abrogated responses relative to WT mice on test day 1, which resulted in a lack of behavioral adaptation between test days 1 and 2 for IP3R2 KO animals (KO – M D1: 0.2 [0-0.5], F D1: 0.3 [0-0.5] ; M D2: 0.2 [0-0.6], F D2: 0.2 [0-0.4]; Fig. 5C). There was no difference between M and F responses for either WT or IP3R2 KO animals. The reduction in responses in IP3R2 KO mice test day 1 was driven by a decrease in “Escape” and “Freeze+escape” behaviors (Fig. S5A-C). While “Escape” behaviors were significantly reduced between WT and IP3R2 KO mice (WT – M D1: 0.4 [0-1.0], D2: 0.4 [0-0.8]; F D1: 0.4 [0-1.0], D2: 0.2 [0-0.5]); KO – M D1: 0.2 [0-0.5], D2: 0 [0-0.6]; F D1: 0.2 [0-0.5], D2: 0.2 [0-0.4]; Fig. S5B), “Freeze+escape” responses were completely abolished in IP3R2 KO mice (WT – M D1: 0.2 [0.0-0.6], D2: 0 [0-0.2]; F D1: 0.2 [0-0.6], D2: 0 [0-0.6]; KO – M D1: 0 [0-0], D2: 0 [0-0]; F D1: 0 [0-0], D1: 0 [0-0]; Fig. S5C). To ensure that these differences were reflecting the responses of the entire cohort, rather than few highly non-responsive individuals, which can skew the average result, we quantified the percent respondent mice (quantified as mice who respond to at least one trial per day) and observed no difference between WT and IP3R2 KO mice across test days for total response (WT – M D1:100% D2: 91% F D1: 100% D2: 73% ; KO – M D1: 91% D2: 55% F D1: 80% D2: 80%) or specific response types (“Freeze” WT – M D1: 36% D2: 18%, F D1: 27% D2: 18%; KO – M D1: 0%, D2: 18%, F D1: 30% D2: 20%; “Escape” WT – M D1: 91% D2: 82%, F D1: 91% D2: 64%; KO – M D1: 91% D2: 36%, F D1: 70% D2: 60%; “Freeze and escape” WT – M D1: 73% D2: 27%, F D1: 64% D2: 64% ; KO – M D1: 0% D2: 0%, F D1: 0% D2: 0%; Fig. 5D, Fig. S5D-F). Importantly, average velocity during the stimulus was not different between the genotypes, suggesting that the observed responses were not due to ambulatory differences (shown as mean ± s.e.m velocity [cm/sec]: WT – M D1: 4.77 ±0.12, D2: 4.60 ±0.19, F D1: 4.86 ±0.19, D2: 4.55 ±0.27; KO – M D1: 5.31 ±0.20, D2: 4.93 ±0.14, F D1: 4.76 ±0.30, D2: 5.07 ±0.29; Fig. 5E). These findings demonstrate that IP3R2 KO mice exhibit an impairment in visually evoked defensive behaviors, despite normal baseline mobility, supporting a role for astrocytic store-released Ca^2+^ signaling in shaping behavior through experience-dependent circuit refinement.

### VC astrocyte morphology is abnormal in IP3R2 knockout mice

We have shown that IP3R2 KO mice VC exhibit gene expression changes relative to WT mice^4^. During development, many changes in astrocytic gene modules affect processes underlying cellular expansion and outgrowth^56^, which is critical for proper synaptic ensheathment in the mature brain^1^. To test whether astrocyte morphology was altered in IP3R2 KO VC, we delivered an AAV via intracerebroventricular (icv) injection to sparsely express the membrane tethered fluorescent reporter (Lck-eGFP) under control of the astrocytic promoter GfaABC_1_D^57^ at P1 and assessed astrocyte morphology two weeks later at P16 (Fig. 6A). The membrane tethered reporter is necessary to fully capture astrocyte morphological complexity, while sparse labeling ensures the ability to analyze individual astrocytes. Lck-eGFP labeling was highly specific (100% of Lck-eGFP expressing cells colocalized with S100β astrocytic marker) and sparse (∼40% of S100ꞵ positive cells expressed Lck-eGFP) confirming the validity of our approach for these experiments (Fig. S6 A-B). 3D volumetric renderings were constructed from the Lck-eGFP signal using Imaris (see Methods) and analyzed for size and geometric qualities (Fig. 6 D-E).

**Figure 6.**
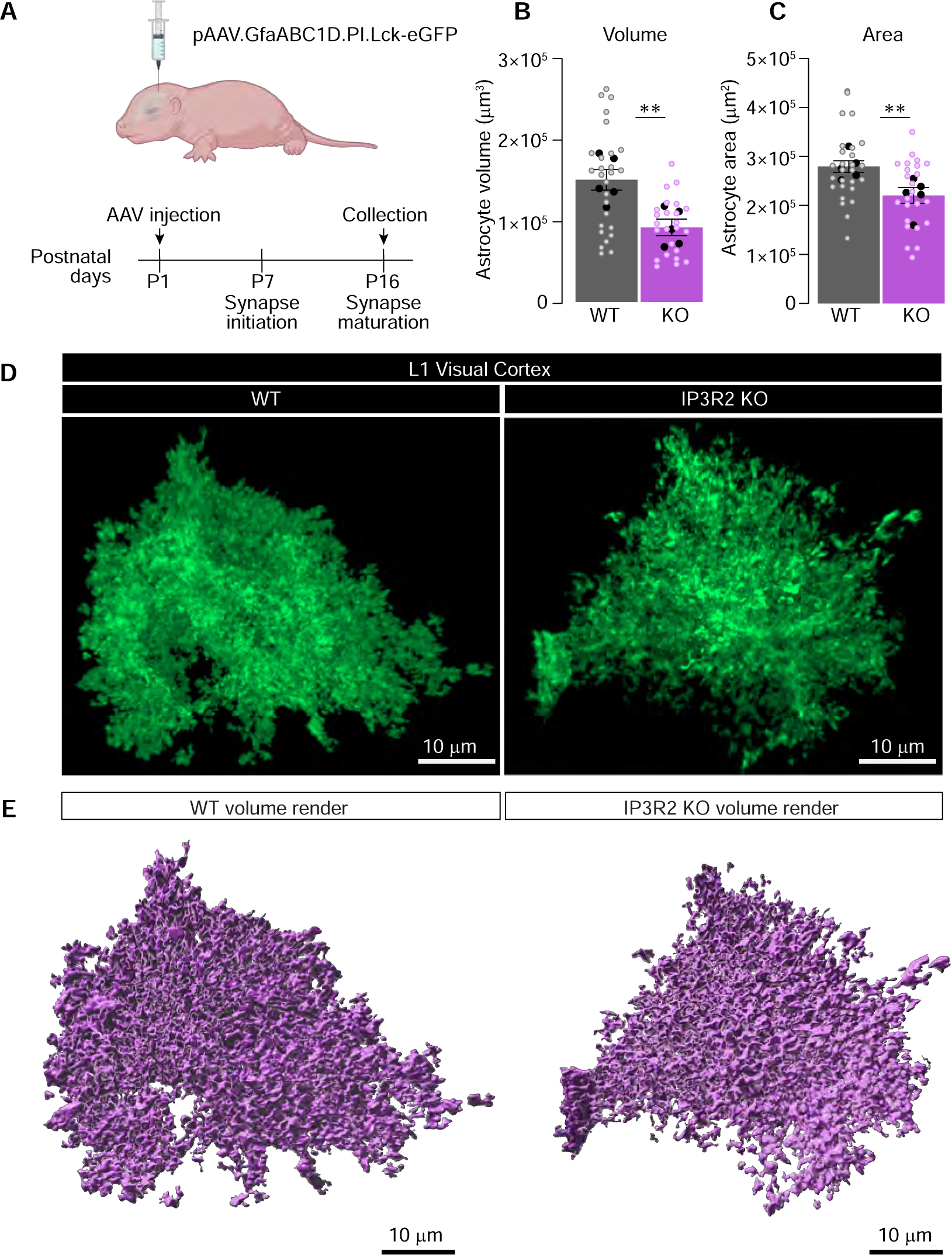
Reduced morphology in IP3R2 KO astrocytes. See also Fig. S6. **A.** Schematic of the strategy to express fluorescent reporter in astrocytes. P1 mouse pups were injected icv with AAV to express membrane tethered eGFP (Lck-eGFP) under astrocyte specific promoter GfaABC1D. Tissue is collected 2 weeks following injection, sectioned and imaged using Airyscan super resolution confocal microscope. **B-E.** Astrocytic volume is reduced in IP3R2 KO mice compared to WT. Example images of Lck-eGFP (green) expressing astrocytes (**D**) and 3D rendering of volume using Imaris (**E**) for each genotype is shown as labeled. **B-C.** Quantification shows reduced total volume and area of IP3R2 KO astrocytes in L1 VC. Graphs show mean ± s.e.m. Black circles above each bar are average of signal in each mouse, colored open circles are data for each astrocyte. Number of mice/ group (N) N=5, number of astrocytes = 25-27. Scale bar = 10 μm. **P<0.01 by t-test.

We observed a significant reduction in total astrocyte volume (∼40%) in IP3R2 KO astrocytes in VC Layer 1 compared to WT (data shown as mean ± s.e.m [µm³]: WT: 14980 ±1270, KO: 9148 ±1011; Fig.6B) and in area (∼20%) (shown as mean ± s.e.m [µm^2^]: WT: 28004 ±1187, KO: 22093 ±1624; Fig. 6C). There were no significant changes in oblate ellipticity (WT: 0.51 ±0.02, KO: 0.54 ±0.03 Fig. S6C), prolate ellipticity (WT: 0.30 ±0.04, KO: 0.28 ±0.04; Fig. S6D), or sphericity (WT: 0.11 ±0.007, KO: 0.10 ±0.005; Fig. S6E) between WT and IP3R2 KO astrocytes. These findings demonstrate that IP3R2 KO astrocytes have morphological deficits compared to WT at P16, a critical developmental timepoint for astrocytic regulation of synapse development. While overall cell size was significantly reduced, the general geometry of IP3R2 KO astrocytes remained consistent with WT astrocytes, suggesting that store-released astrocytic Ca^2+^ signaling modulates astrocyte outgrowth while fundamental cytoskeletal and membrane patterning mechanisms are likely preserved.

## DISCUSSION

This study reveals that astrocytic IP3R2 mediated Ca^2+^ signaling is essential for the maturation and refinement of excitatory synapses in the developing visual cortex. Specifically, we show:

- IP3R2 KO mice exhibit reduced numbers and volumes of presynaptic VGLUTs and VGLUT-containing synapses at P14 and P28, but not P7. These reductions are specific to glutamatergic presynaptic terminals, with no changes in axonal innervation to the visual cortex, total cell number, or inhibitory GABAergic synapses.
- Visually evoked neuronal c-FOS activation in response to a light pulse is reduced in IP3R2 KO mice across visual circuit brain regions, but baseline levels of neuronal c-FOS activity remian unaffected.
- Defensive behavioral responses to a looming stimulus are blunted in IP3R2 KO mice.
- Astrocyte volume and area are decreased in the VC of IP3R2 KO mice at P16 during the period of peak synaptogenesis.

### Astrocytic ER store-released Ca^2+^ supports synaptic development

The appropriate spatio-temporal expression of synaptic proteins is critical for proper synapse maturation in the developing brain. In IP3R2 KO mice, we observed a developmentally specific reduction of presynaptic VGLUTs (Fig. 1, Fig. S1, Fig. S2). Accompanied by our finding that IP3R2 protein levels were stable across development (Fig. 1B), this suggests that IP3R2 signaling becomes increasingly important as synapses mature. While initial stages of synaptogenesis in the VC may be driven by neuron autonomous programs, later timepoints, such as the period following eye opening, likely require astrocytic input mediated by store-released Ca^2+^ signaling to stabilize functional connections. This aligns with previous work showing that astrocytic gene expression becomes increasingly specialized during specific stages of synapse development, and that Ca^2+^-dependent transcriptional regulation is a major mechanism by which astrocytes influence circuit assembly^4,12^. Astrocytic store-released Ca^2+^ from the ER plays a central role in coordinating astrocytic responses to neuronal activity^24,31^, and our data suggest that this signaling axis is required for the proper maturation of excitatory terminals.

Importantly, synaptic deficits in IP3R2 KO mice were limited to glutamatergic circuits, with inhibitory synapses remaining intact (Fig. 3). This synapse type specific effect has important implications for cortical excitatory/inhibitory (E/I) balance. Disruptions in E/I ratio are a hallmark of multiple neurological disorders^58–61^, and previous studies have linked astrocyte Ca^2+^ signaling to their pathogenesis. For example, it was shown that IP3R2 KO mice exhibit disrupted resting-state functional connectivity in medial prefrontal cortex (mPFC)-centered networks, mirroring patterns observed in humans with major depressive disorder^62^, which is characterized by disrupted glutamate and GABA signaling^6^. Early life disruptions in glutamatergic synapses caused by dysregulated astrocytic store-released Ca^2+^ signaling could precede the development of these disorders in the mature brain. Ongoing research focusing on identifying the downstream Ca^2+^ dependent pathways in astrocytes that selectively influence excitatory synapse maturation and maintain E/I balance during critical developmental periods will be essential to our understanding of these functions.

### IP3R2 mediated Ca^2+^ signaling contributes to the development of visual circuit function and behavioral output

At the circuit level, we found that evoked (but not basal) neuronal activation was impaired in IP3R2 KO mice. Expression of the immediate early gene c-FOS was significantly reduced in response to visual stimulation across both cortical and subcortical structures (Fig. 4, Fig. S4). These findings are consistent with previous reports demonstrating that astrocytic Ca^2+^ transients contribute to stimulus dependent neuronal activation and cortical state transitions, particularly in sensory cortices where astrocyte Ca^2+^ activity has been shown to gate sensory throughput and regulate network gain^44,63^. Specifically, IP3R2 KO was shown to elevate sensory evoked gamma activity, an effect that reflects reduced astrocyte-mediated modulation of cortical excitability and aligns with our observation of impaired neuronal activation following visual stimulation. Further, the suppression of visually evoked c-FOS expression in IP3R2 KO mice is consistent with work showing that astrocyte mediated modulation of excitatory neurotransmission is critical for experience-dependent plasticity in early postnatal visual circuits^64^.

At the behavioral level, these circuit impairments translated into abrogated responses to looming visual threats (Fig. 5, Fig. S5). These phenotypes align with reports detailing disrupted social behavior in adult IP3R2 KO mice, specifically highlighting temporal delays in dominance behaviors and impaired social interaction^22,31^. Because social behaviors rely in part on acute visual system activation and integration, deficits in excitatory circuit maturation which lead to reductions in context dependent neuronal activation in IP3R2 KO mice could underlie these effects.

In contrast, several studies reported that mice lacking IP3R2 have no discernable changes in synaptic activity or behavior^65,66^ These discrepancies could be due to regional specificity (e.g. different regulatory pathways in the hippocampus, where no changes in synaptic activity were observed, compared to the cortex), developmental stage, or in the types of behavioral assessments. For example, it was shown^31^ that the *onset latency* of social dominance behavior, but not the behavior itself, was disrupted in IP3R2 KO mice. In accordance, we observed that unique components of defensive behaviors were differentially modulated in IP3R2 KO mice (Fig. S5). Thus, while astrocytic store-released Ca^2+^ probably is not responsible for every aspect of behavioral outputs that it is involved in, it likely plays important roles in shaping their specific components or temporal scales, an idea that aligns with recent reports implying that astrocytic Ca^2+^ is at the interface of behavioral state transitions^67–70^. Additionally, some astrocytic Ca^2+^ linked behaviors occur entirely independent of IP3R2 mediated signaling^25^, emphasizing a need for studies that quantify how different types of astrocytic Ca^2+^ signals integrate to produce functional outputs.

These findings collectively underscore the importance of considering regional specificity, Ca^2+^ source interactions, and modular components of behavioral outputs in studies investing the effects of IP3R2 mediated Ca^2+^ signaling in neuronal network modulation and behavioral responses. They also suggest that this signaling pathway plays an important role in these processes by modulating neuronal responsiveness to salient stimuli, consistent with studies implicating astrocyte Ca^2+^ in encoding behavioral salience and coordinating network level responses during exploratory and defensive states^18,25,71^. Future studies will be necessary to determine how astrocytic Ca^2+^ signaling interfaces with neuromodulatory systems and whether restoring evoked responses in IP3R2 deficient circuits can rescue behavioral deficits.

### Store-Released Ca^2+^ Regulates Astrocyte Morphological Development

A major question that remains involves the potential mechanisms by which astrocytes may induce their regulatory effects on synaptic maturation, and the role that store-released Ca^2+^ plays in this function. We approached this question by quantifying morphology in WT and IP3R2 KO mice, demonstrating that mice lacking IP3R2 have reduced astrocyte volume and area in the visual cortex, suggesting a possible mechanistic link. This aligns with studies demonstrating that Ca^2+^ dependent pathways are relevant to astrocyte growth and structural maturation^56,72^. Appropriate outgrowth and process elaboration is critical for proper synaptic ensheathment and gliotransmitter release, and loss or reduction of these processes could underlie deficits in astrocyte-neuron signaling, thus impairing synaptic integrity. Interestingly, while overall volume and surface area were reduced in IP3R2 KOs, the geometric parameters of astrocyte shape such as sphericity and ellipticity were preserved. This suggests that IP3R2-mediated Ca^2+^ signaling selectively regulates growth rather than global cytoskeletal patterning mechanisms. Further investigation is needed to determine whether these morphological deficits disrupt functional astrocyte–synapse interactions *in vivo* and how they relate to gliotransmission or synapse specificity.

In summary, in the present study we demonstrate that IP3R2 mediated astrocytic Ca^2+^ signaling is an essential intermediary in the development and function of excitatory synapses in the mouse visual system. However, astrocytic Ca^2+^ signaling is highly heterogeneous spatially, temporally, and mechanistically^15,26,27,29,42,73–75^. Our work highlights the role of one defined signaling pathway (ER store-released Ca^2+^ via IP3R2) in a specific brain region, and in a developmental context. These findings underscore the need to better understand how astrocytes integrate diverse Ca^2+^ signals to coordinate their functions across circuits. To fully appreciate and comprehend these complex relationships, studies combining astrocyte specific manipulations, *in vivo* imaging, and computational modeling will be essential to our understanding of how distinct astrocytic Ca^2+^ signals contribute to synapse development, circuit function, and behavior.

## RESOURCE AVAILABILITY

Requests for further information and resources should be directed to and will be fulfilled by the lead contact, Isabella Farhy-Tselnicker.

## ACKNOWLEDGEMENTS

This work was funded by NIH R01NS133047 to IFT, and NS133047 to JM.

## AUTHOR CONTRIBUTIONS

GI conceptualized, conducted experiments, and performed data analysis. JM performed the behavioral experiments. MG performed WB experiments. IFT conceived the project, supervised experiments and data analysis, and wrote the paper with GI and input from other authors.

## SUPPLEMENTAL INFORMATION

Document S1. 5 Supplemental figures. Figures S1, S2, S4, S5, S6.

## REFERENCES

1 Chalmers, N., Masouti, E. & Beckervordersandforth, R. Astrocytes in the adult dentate gyrus-balance between adult and developmental tasks. Mol Psychiatry 29, 982–991 (2024). 10.1038/s41380-023-02386-4

2 Farhy-Tselnicker, I. & Allen, N. J. Astrocytes, neurons, synapses: a tripartite view on cortical circuit development. Neural Development 13, 7 (2018). 10.1186/s13064-018-0104-y

3 Tan, C. X., Burrus Lane, C. J. & Eroglu, C. Role of astrocytes in synapse formation and maturation. Curr Top Dev Biol 142, 371–407 (2021). 10.1016/bs.ctdb.2020.12.010

4 Farhy-Tselnicker, I. et al. Activity-dependent modulation of synapse-regulating genes in astrocytes. Elife 10 (2021). 10.7554/eLife.70514

5 Xie, Y. et al. Astrocyte-neuron crosstalk through Hedgehog signaling mediates cortical synapse development. Cell Rep 38, 110416 (2022). 10.1016/j.celrep.2022.110416

6 Duman, R. S., Sanacora, G. & Krystal, J. H. Altered Connectivity in Depression: GABA and Glutamate Neurotransmitter Deficits and Reversal by Novel Treatments. Neuron 102, 75–90 (2019). 10.1016/j.neuron.2019.03.013

7 Guang, S. et al. Synaptopathology Involved in Autism Spectrum Disorder. Front Cell Neurosci 12, 470 (2018). 10.3389/fncel.2018.00470

8. Bernard, C. Jasper’s Basic Mechanisms of the Epilepsies (eds J. L. Noebels et al.) (2012).

9 Li, H. et al. Laminar and columnar development of barrel cortex relies on thalamocortical neurotransmission. Neuron 79, 970–986 (2013). 10.1016/j.neuron.2013.06.043

10 Blue, M. E. & Parnavelas, J. G. The formation and maturation of synapses in the visual cortex of the rat. II. Quantitative analysis. J Neurocytol 12, 697–712 (1983). 10.1007/BF01181531

11 Boisvert, M. M., Erikson, G. A., Shokhirev, M. N. & Allen, N. J. The Aging Astrocyte Transcriptome from Multiple Regions of the Mouse Brain. Cell Rep 22, 269–285 (2018). 10.1016/j.celrep.2017.12.039

12 Imrie, G., Gray, M. B., Raghuraman, V. & Farhy-Tselnicker, I. Gene Expression at the Tripartite Synapse: Bridging the Gap Between Neurons and Astrocytes. Adv Neurobiol 39, 95–136 (2024). 10.1007/978-3-031-64839-7_5

13 Faust, T. E. et al. Glial Control of Cortical Neuronal Circuit Maturation and Plasticity. J Neurosci 44 (2024). 10.1523/JNEUROSCI.1208-24.2024

14 Adamsky, A. et al. Astrocytic Activation Generates De Novo Neuronal Potentiation and Memory Enhancement. Cell 174, 59–71 e14 (2018). 10.1016/j.cell.2018.05.002

15 Ahrens, M. B., Khakh, B. S. & Poskanzer, K. E. Astrocyte Calcium Signaling. Cold Spring Harb Perspect Biol 16 (2024). 10.1101/cshperspect.a041353

16 Bai, Y. et al. Revisiting astrocytic calcium signaling in the brain. Fundam Res 4, 1365–1374 (2024). 10.1016/j.fmre.2023.11.021

17 Imrie, G. & Farhy-Tselnicker, I. Astrocyte regulation of behavioral outputs: the versatile roles of calcium. Front Cell Neurosci 19, 1606265 (2025). 10.3389/fncel.2025.1606265

18 Cho, W. H. et al. Hippocampal astrocytes modulate anxiety-like behavior. Nat Commun 13, 6536 (2022). 10.1038/s41467-022-34201-z

19 Peyton, L. et al. In vivo calcium extrusion from accumbal astrocytes reduces anxiety-like behaviors but increases compulsive-like responses and compulsive ethanol drinking in mice. Neuropharmacology 268, 110320 (2025). 10.1016/j.neuropharm.2025.110320

20 Rupprecht, P. et al. Centripetal integration of past events in hippocampal astrocytes regulated by locus coeruleus. Nat Neurosci 27, 927–939 (2024). 10.1038/s41593-024-01612-8

21 Suthard, R. L. et al. Chronic Gq activation of ventral hippocampal neurons and astrocytes differentially affects memory and behavior. Neurobiol Aging 125, 9–31 (2023). 10.1016/j.neurobiolaging.2023.01.007

22 Wang, Q. et al. Impaired calcium signaling in astrocytes modulates autism spectrum disorder-like behaviors in mice. Nat Commun 12, 3321 (2021). 10.1038/s41467-021-23843-0

23 Yu, X. et al. Reducing Astrocyte Calcium Signaling In Vivo Alters Striatal Microcircuits and Causes Repetitive Behavior. Neuron 99, 1170–1187 e1179 (2018). 10.1016/j.neuron.2018.08.015

24 Sherwood, M. W., Arizono, M., Panatier, A., Mikoshiba, K. & Oliet, S. H. R. Astrocytic IP(3)Rs: Beyond IP(3)R2. Front Cell Neurosci 15, 695817 (2021). 10.3389/fncel.2021.695817

25 Srinivasan, R. et al. Ca(2+) signaling in astrocytes from Ip3r2(-/-) mice in brain slices and during startle responses in vivo. Nat Neurosci 18, 708–717 (2015). 10.1038/nn.4001

26 Agarwal, A. et al. Transient Opening of the Mitochondrial Permeability Transition Pore Induces Microdomain Calcium Transients in Astrocyte Processes. Neuron 93, 587–605 e587 (2017). 10.1016/j.neuron.2016.12.034

27 Ahmadpour, N., Kantroo, M. & Stobart, J. L. Extracellular Calcium Influx Pathways in Astrocyte Calcium Microdomain Physiology. Biomolecules 11 (2021). 10.3390/biom11101467

28 Denizot, A., Arizono, M., Nagerl, U. V., Berry, H. & De Schutter, E. Control of Ca(2+) signals by astrocyte nanoscale morphology at tripartite synapses. Glia 70, 2378–2391 (2022). 10.1002/glia.24258

29 Lia, A. et al. Calcium Signals in Astrocyte Microdomains, a Decade of Great Advances. Front Cell Neurosci 15, 673433 (2021). 10.3389/fncel.2021.673433

30 Shigetomi, E., Jackson-Weaver, O., Huckstepp, R. T., O’Dell, T. J. & Khakh, B. S. TRPA1 channels are regulators of astrocyte basal calcium levels and long-term potentiation via constitutive D-serine release. J Neurosci 33, 10143–10153 (2013). 10.1523/JNEUROSCI.5779-12.2013

31 Guillot de Suduiraut, I., Grosse, J., Ramos-Fernandez, E., Sandi, C. & Hollis, F. Astrocytic release of ATP through type 2 inositol 1,4,5-trisphosphate receptor calcium signaling and social dominance behavior in mice. Eur J Neurosci 53, 2973–2985 (2021). 10.1111/ejn.14892

32 Jego, P., Pacheco-Torres, J., Araque, A. & Canals, S. Functional MRI in mice lacking IP3-dependent calcium signaling in astrocytes. J Cereb Blood Flow Metab 34, 1599–1603 (2014). 10.1038/jcbfm.2014.144

33 Li, X., Zima, A. V., Sheikh, F., Blatter, L. A. & Chen, J. Endothelin-1-induced arrhythmogenic Ca2+ signaling is abolished in atrial myocytes of inositol-1,4,5-trisphosphate(IP3)-receptor type 2-deficient mice. Circ Res 96, 1274–1281 (2005). 10.1161/01.RES.0000172556.05576.4c

34 Glascock, J. J. et al. Delivery of therapeutic agents through intracerebroventricular (ICV) and intravenous (IV) injection in mice. J Vis Exp (2011). 10.3791/2968

35 Kawasaki, H. et al. Intracerebroventricular and Intravascular Injection of Viral Particles and Fluorescent Microbeads into the Neonatal Brain. J Vis Exp (2016). 10.3791/54164

36 Kim, J. Y., Grunke, S. D., Levites, Y., Golde, T. E. & Jankowsky, J. L. Intracerebroventricular viral injection of the neonatal mouse brain for persistent and widespread neuronal transduction. J Vis Exp, 51863 (2014). 10.3791/51863

37 Daviu, N. et al. Visual-looming Shadow Task with in-vivo Calcium Activity Monitoring to Assess Defensive Behaviors in Mice. Bio Protoc 10, e3826 (2020). 10.21769/BioProtoc.3826

38 Daviu, N. et al. Paraventricular nucleus CRH neurons encode stress controllability and regulate defensive behavior selection. Nat Neurosci 23, 398–410 (2020). 10.1038/s41593-020-0591-0

39 Mathis, A. et al. DeepLabCut: markerless pose estimation of user-defined body parts with deep learning. Nat Neurosci 21, 1281–1289 (2018). 10.1038/s41593-018-0209-y

40 Nath, T. et al. Using DeepLabCut for 3D markerless pose estimation across species and behaviors. Nat Protoc 14, 2152–2176 (2019). 10.1038/s41596-019-0176-0

41 Gabriel, C. J. et al. BehaviorDEPOT is a simple, flexible tool for automated behavioral detection based on markerless pose tracking. Elife 11 (2022). 10.7554/eLife.74314

42 Khakh, B. S. & McCarthy, K. D. Astrocyte calcium signaling: from observations to functions and the challenges therein. Cold Spring Harb Perspect Biol 7, a020404 (2015). 10.1101/cshperspect.a020404

43 Veiga, A. et al. Calcium-Dependent Signaling in Astrocytes: Downstream Mechanisms and Implications for Cognition. J Neurochem 169, e70019 (2025). 10.1111/jnc.70019

44 Lines, J., Martin, E. D., Kofuji, P., Aguilar, J. & Araque, A. Astrocytes modulate sensory-evoked neuronal network activity. Nat Commun 11, 3689 (2020). 10.1038/s41467-020-17536-3

45 Zhi, J. J. et al. Insufficient Oligodendrocyte Turnover in Optic Nerve Contributes to Age-Related Axon Loss and Visual Deficits. J Neurosci 43, 1859–1870 (2023). 10.1523/JNEUROSCI.2130-22.2023

46 Benn, S. C., Costigan, M., Tate, S., Fitzgerald, M. & Woolf, C. J. Developmental expression of the TTX-resistant voltage-gated sodium channels Nav1.8 (SNS) and Nav1.9 (SNS2) in primary sensory neurons. J Neurosci 21, 6077–6085 (2001). 10.1523/JNEUROSCI.21-16-06077.2001

47 Sawant, L. A., Hasgekar, N. N. & Vyasarayani, L. S. Developmental expression of neurofilament and glial filament proteins in rat cerebellum. Int J Dev Biol 38, 429–437 (1994).

48 Wei, J. R. et al. Identification of visual cortex cell types and species differences using single-cell RNA sequencing. Nat Commun 13, 6902 (2022). 10.1038/s41467-022-34590-1

49 Ellis, E. M., Gauvain, G., Sivyer, B. & Murphy, G. J. Shared and distinct retinal input to the mouse superior colliculus and dorsal lateral geniculate nucleus. J Neurophysiol 116, 602–610 (2016). 10.1152/jn.00227.2016

50 Schiapparelli, L. M. et al. The Retinal Ganglion Cell Transportome Identifies Proteins Transported to Axons and Presynaptic Compartments in the Visual System In Vivo. Cell Rep 28, 1935–1947 e1935 (2019). 10.1016/j.celrep.2019.07.037

51 Ahmadlou, M., Zweifel, L. S. & Heimel, J. A. Functional modulation of primary visual cortex by the superior colliculus in the mouse. Nat Commun 9, 3895 (2018). 10.1038/s41467-018-06389-6

52 White, B. J., Kan, J. Y., Levy, R., Itti, L. & Munoz, D. P. Superior colliculus encodes visual saliency before the primary visual cortex. Proc Natl Acad Sci U S A 114, 9451–9456 (2017). 10.1073/pnas.1701003114

53 Narushima, M., Agetsuma, M. & Nabekura, J. Development and experience-dependent modulation of the defensive behaviors of mice to visual threats. J Physiol Sci 72, 5 (2022). 10.1186/s12576-022-00831-7

54 Shamash, P. & Branco, T. Protocol to Study Spatial Subgoal Learning Using Escape Behavior in Mice. Bio Protoc 12, e4443 (2022). 10.21769/BioProtoc.4443

55 Vale, R., Evans, D. A. & Branco, T. Rapid Spatial Learning Controls Instinctive Defensive Behavior in Mice. Curr Biol 27, 1342–1349 (2017). 10.1016/j.cub.2017.03.031

56 Chen, J. et al. Astrocyte growth is driven by the Tre1/S1pr1 phospholipid-binding G protein-coupled receptor. Neuron 112, 93–112 e110 (2024). 10.1016/j.neuron.2023.11.008

57 Wu, L. et al. The cell-surface shared proteome of astrocytes and neurons and the molecular foundations of their multicellular interactions. Neuron (2025). 10.1016/j.neuron.2025.05.019

58 Hu, Y. T., Tan, Z. L., Hirjak, D. & Northoff, G. Brain-wide changes in excitation-inhibition balance of major depressive disorder: a systematic review of topographic patterns of GABA- and glutamatergic alterations. Mol Psychiatry 28, 3257–3266 (2023). 10.1038/s41380-023-02193-x

59 Li, G. et al. Revealing excitation-inhibition imbalance in Alzheimer’s disease using multiscale neural model inversion of resting-state functional MRI. Commun Med (Lond) 5, 17 (2025). 10.1038/s43856-025-00736-7

60 Sylvester, A. L. et al. Neural excitation/inhibition imbalance and neurodevelopmental pathology in human copy number variant syndromes: a systematic review. J Neurodev Disord 17, 31 (2025). 10.1186/s11689-025-09614-8

61 Liu, Y. et al. A Selective Review of the Excitatory-Inhibitory Imbalance in Schizophrenia: Underlying Biology, Genetics, Microcircuits, and Symptoms. Front Cell Dev Biol 9, 664535 (2021). 10.3389/fcell.2021.664535

62 Liu, J. et al. Astrocyte dysfunction drives abnormal resting-state functional connectivity in depression. Sci Adv 8, eabo2098 (2022). 10.1126/sciadv.abo2098

63 Cahill, M. K. et al. Network-level encoding of local neurotransmitters in cortical astrocytes. Nature 629, 146–153 (2024). 10.1038/s41586-024-07311-5

64 Reitman, M. E. et al. Norepinephrine links astrocytic activity to regulation of cortical state. Nat Neurosci 26, 579–593 (2023). 10.1038/s41593-023-01284-w

65 Petravicz, J., Boyt, K. M. & McCarthy, K. D. Astrocyte IP3R2-dependent Ca(2+) signaling is not a major modulator of neuronal pathways governing behavior. Front Behav Neurosci 8, 384 (2014). 10.3389/fnbeh.2014.00384

66 Petravicz, J., Fiacco, T. A. & McCarthy, K. D. Loss of IP3 receptor-dependent Ca2+ increases in hippocampal astrocytes does not affect baseline CA1 pyramidal neuron synaptic activity. J Neurosci 28, 4967–4973 (2008). 10.1523/JNEUROSCI.5572-07.2008

67 Oliveira, J. F. & Araque, A. Astrocyte regulation of neural circuit activity and network states. Glia 70, 1455–1466 (2022). 10.1002/glia.24178

68 Bojarskaite, L. et al. Astrocytic Ca(2+) signaling is reduced during sleep and is involved in the regulation of slow wave sleep. Nat Commun 11, 3240 (2020). 10.1038/s41467-020-17062-2

69 Peng, W. et al. Adenosine-independent regulation of the sleep-wake cycle by astrocyte activity. Cell Discov 9, 16 (2023). 10.1038/s41421-022-00498-9

70 Wang, F. et al. Distinct astrocytic modulatory roles in sensory transmission during sleep, wakefulness, and arousal states in freely moving mice. Nat Commun 14, 2186 (2023). 10.1038/s41467-023-37974-z

71 Gau, Y. A. et al. Multicore fiber optic imaging reveals that astrocyte calcium activity in the mouse cerebral cortex is modulated by internal motivational state. Nat Commun 15, 3039 (2024). 10.1038/s41467-024-47345-x

72 Bernardinelli, Y. et al. Activity-dependent structural plasticity of perisynaptic astrocytic domains promotes excitatory synapse stability. Curr Biol 24, 1679–1688 (2014). 10.1016/j.cub.2014.06.025

73 Oberheim, N. A., Goldman, S. A. & Nedergaard, M. Heterogeneity of astrocytic form and function. Methods Mol Biol 814, 23–45 (2012). 10.1007/978-1-61779-452-0_3

74 Bindocci, E. et al. Three-dimensional Ca(2+) imaging advances understanding of astrocyte biology. Science 356 (2017). 10.1126/science.aai8185

75 Bosson, A. et al. TRPA1 channels promote astrocytic Ca(2+) hyperactivity and synaptic dysfunction mediated by oligomeric forms of amyloid-beta peptide. Mol Neurodegener 12, 53 (2017). 10.1186/s13024-017-0194-8

