## Supplemental Document S1 for "Astrocyte Store-Released Calcium Modulates Visual Cortex Synapse Development and Circuit Function"

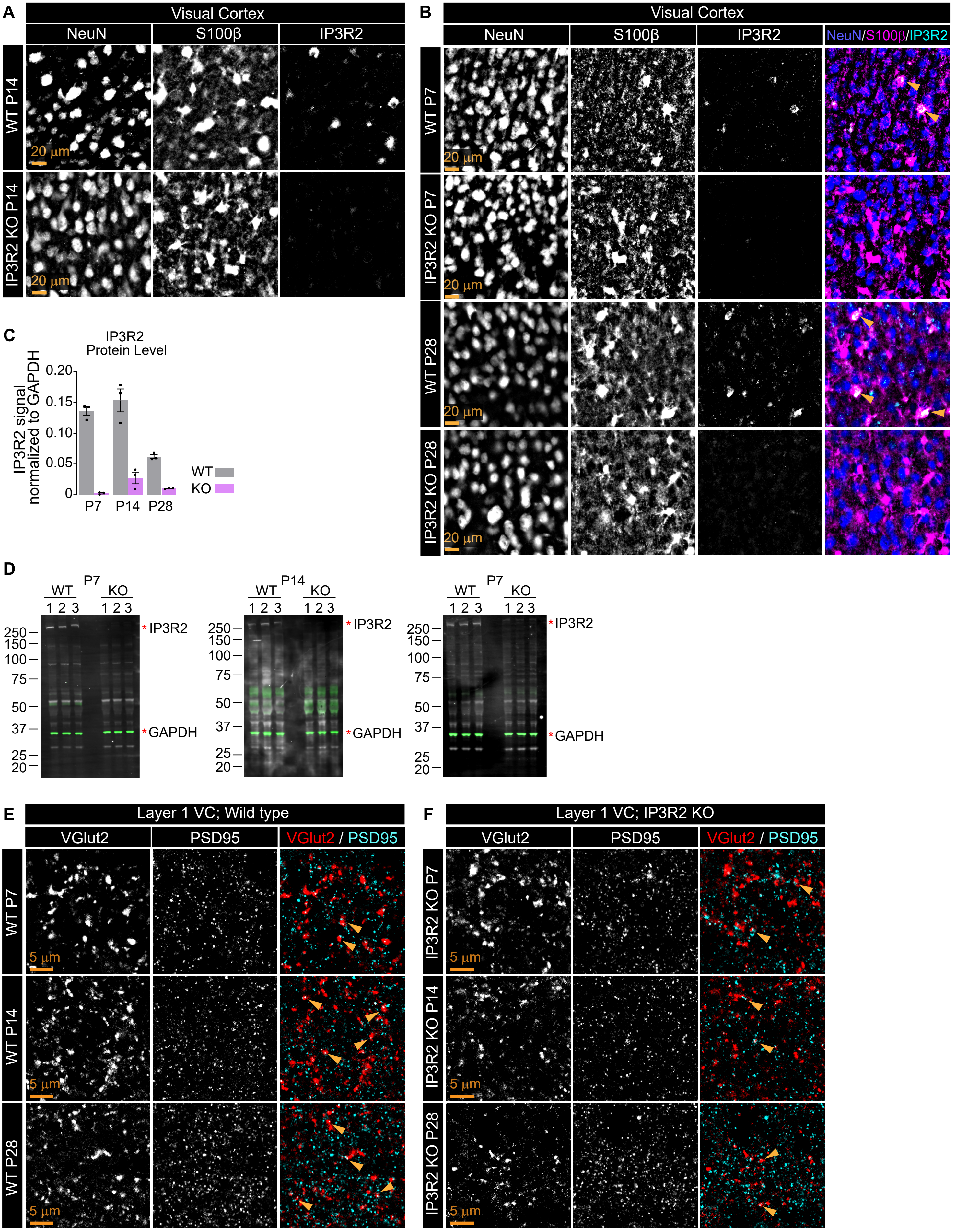
A
Visual Cortex
NeuN
S100β
IP3R2
WT P14
IP3R2 KO P14
20 µm
B
Visual Cortex
NeuN
S100β
IP3R2
NeuN/S100β/IP3R2
WT P7
IP3R2 KO P7
WT P28
IP3R2 KO P28
20 µm
C
IP3R2 Protein Level
IP3R2 signal normalized to GAPDH
0.20
0.15
0.10
0.05
0
P7
P14
P28
WT
KO
D
P7
P14
P7
WT
KO
1 2 3
250
150
100
75
50
37
25
20
*IP3R2
*GAPDH
E
Layer 1 VC; Wild type
VGlut2
PSD95
VGlut2 / PSD95
WT P7
WT P14
WT P28
5 µm
F
Layer 1 VC; IP3R2 KO
VGlut2
PSD95
VGlut2 / PSD95
IP3R2 KO P7
IP3R2 KO P14
IP3R2 KO P28
5 µm

**Figure S1. Glutamatergic synapse development is perturbed in IP3R2 KO mice VC. A.** Example single channel images of P14 VC in WT and KO mice as labeled showing Neun, S100β, and IP3R2 signal. **B.** Example images of Neun, S100β and IP3R2 in WT and KO VC at P7 and P28 as labeled. Single channel images are on the left, merged images on the right. **C.** Quantification of Western Blots represented as IP3R2 signal normalized to GAPDH loading control for each age and genotype as labeled. IP3R2 signal is strongly reduced in IP3R2 KO mice. **D.** Uncropped Western Blot membranes showing IP3R2 signal (white; imaged at 700 channel) and GAPDH signal (green imaged at 800 channel) for each age and genotype as labeled. Numbers indicate individual animal samples. **E-F.** Example images of the presynaptic VGLUT2, postsynaptic PSD95 and merged (synapses) in each age and genotype as labeled. Single channel images on the left, merged images on the right.

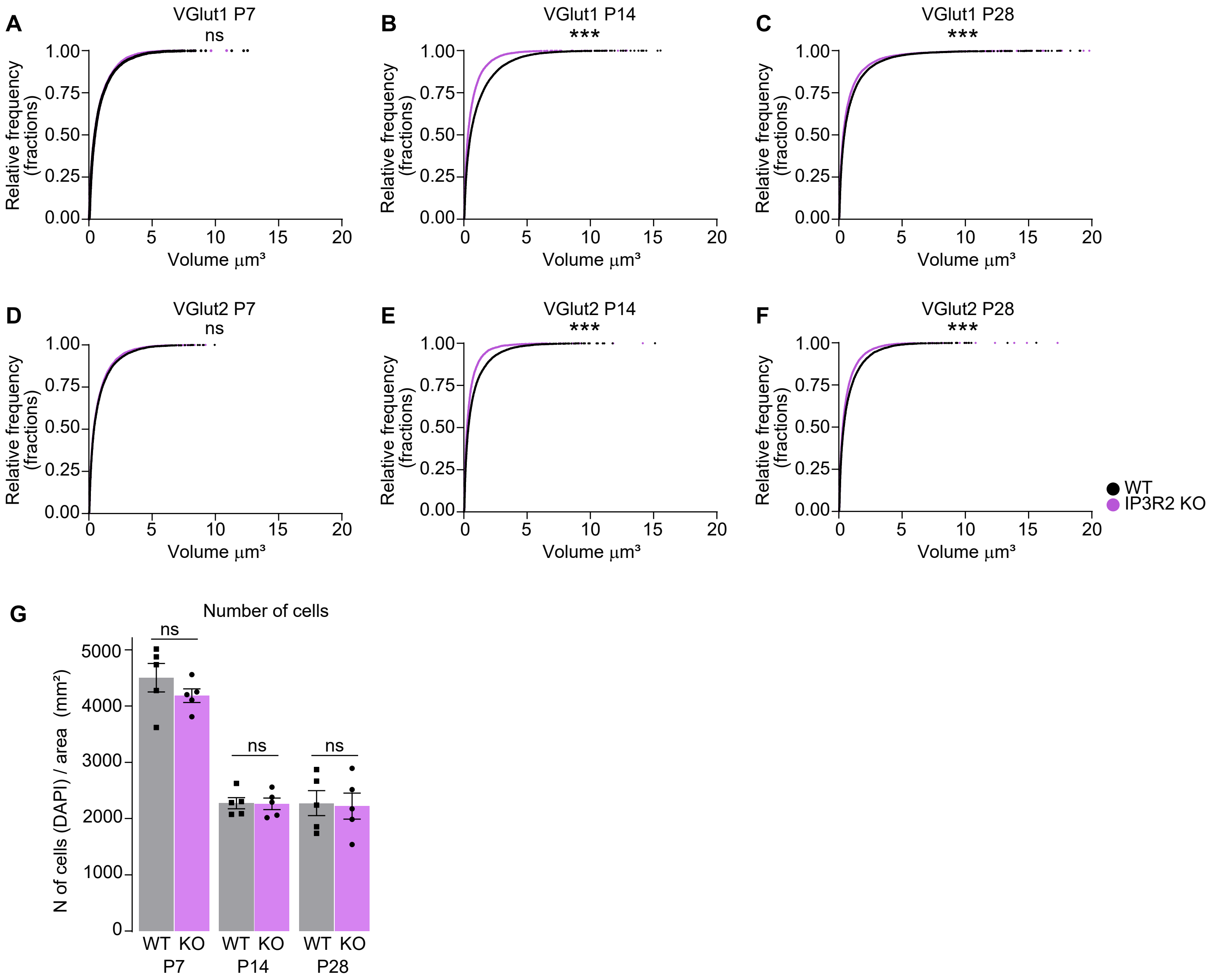

A
VGlut1 P7
ns
B
VGlut1 P14
***
C
VGlut1 P28
***
D
VGlut2 P7
ns
E
VGlut2 P14
***
F
VGlut2 P28
***
Relative frequency (fractions)
1.00
0.75
0.50
0.25
0.00
Volume μm³
0
5
10
15
20
WT
IP3R2 KO
G
Number of cells
N of cells (DAPI) / area (mm²)
5000
4000
3000
2000
1000
0
ns
WT KO
P7
P14
P28

**Figure S2. VGLUT volumes are reduced in IP3R2 KO mice with no change in cell numbers. A-F.** Cumulative distributions of volumes from 3D rendered images (see Fig. 1, Fig. S1) for VGLUT1 (**A-C**) and VGLUT2 (**D-F**) per age and genotype as labeled. Higher fraction of smaller volumes (<5 $\mu m^3$) is observed in IP3R2 KO mice at P14 and P28, but not P7. ***P<0.001 by Kolmogorov-Smirnov test comparing WT and KO within each age. ns denotes non-significant results (>0.05). **G.** Number of cells is unaltered by IP3R2 KO at any age tested. Graph shows mean ± s.e.m. Squares and circles above each bar are average of signal in each mouse. Quantification was done from images in Fig. 2A-B. Number of mice/ group (N) N=5.

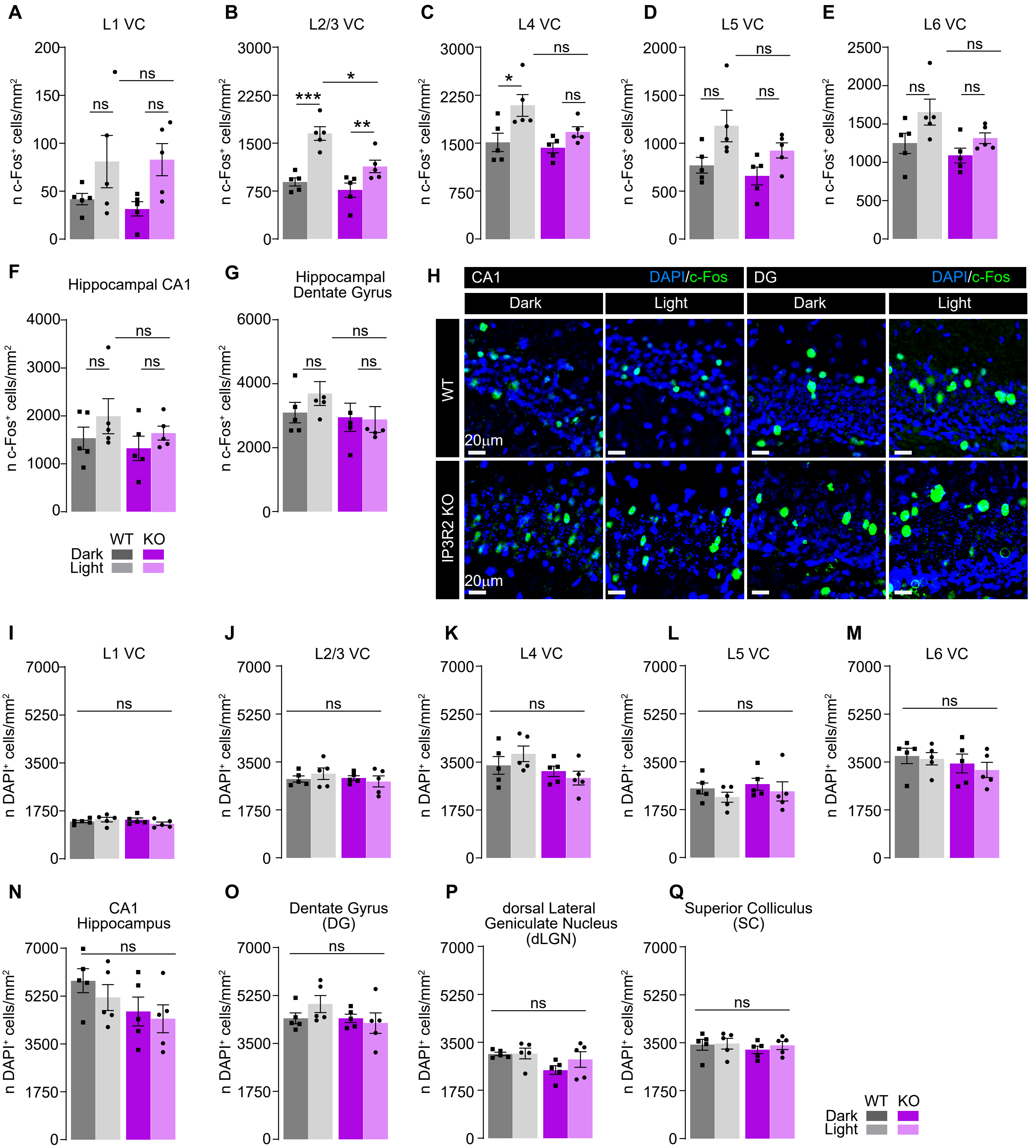

A
L1 VC
B
L2/3 VC
C
L4 VC
D
L5 VC
E
L6 VC
n c-Fos⁺ cells/mm²
F
Hippocampal CA1
G
Hippocampal Dentate Gyrus
H
CA1
DAPI/c-Fos
DG
DAPI/c-Fos
Dark
Light
Dark
Light
WT
IP3R2 KO
20μm
WT KO
Dark
Light
I
L1 VC
J
L2/3 VC
K
L4 VC
L
L5 VC
M
L6 VC
n DAPI⁺ cells/mm²
N
CA1 Hippocampus
O
Dentate Gyrus (DG)
P
dorsal Lateral Geniculate Nucleus (dLGN)
Q
Superior Colliculus (SC)
WT KO
Dark
Light

**Figure S4. Light-evoked c-FOS expression is blunted in IP3R2 KO mice visual circuit. A-E.** Light evoked c-FOS levels are diminished in IP3R2 KO mice within specific layers of the VC. Quantification is represented as c-FOS positive cell numbers per area. Light exposure produced a significant increase in the number of c-FOS positive cells in the WT only in layers 2-3 and 4, but not in the KO. **F-H.** Light pulse does not induce c-FOS expression in the hippocampus. Example images of c-FOS (green) and DAPI to mark cell nuclei (blue) in the CA1 and DG regions of the hippocampus in both genotypes as labeled. Quantification of CA1 (**F**) and DG (**G**) is also shown. No difference is observed between light and dark conditions or the genotypes in either sub-region. **I-Q.** Quantification of DAPI positive cells shows light induced c-FOS increase is not due to increased cell number, and no difference between genotypes is observed. Graphs show mean ± s.e.m. Squares and circles above each bar are average of signal in each mouse. Number of mice/ group (N) N=5, KO DG N=4. Scale bar = 20 μm. *P<0.05, **P<0.01, ***P<0.001 by one-way ANOVA. ns denotes non-significant results (P>0.05).

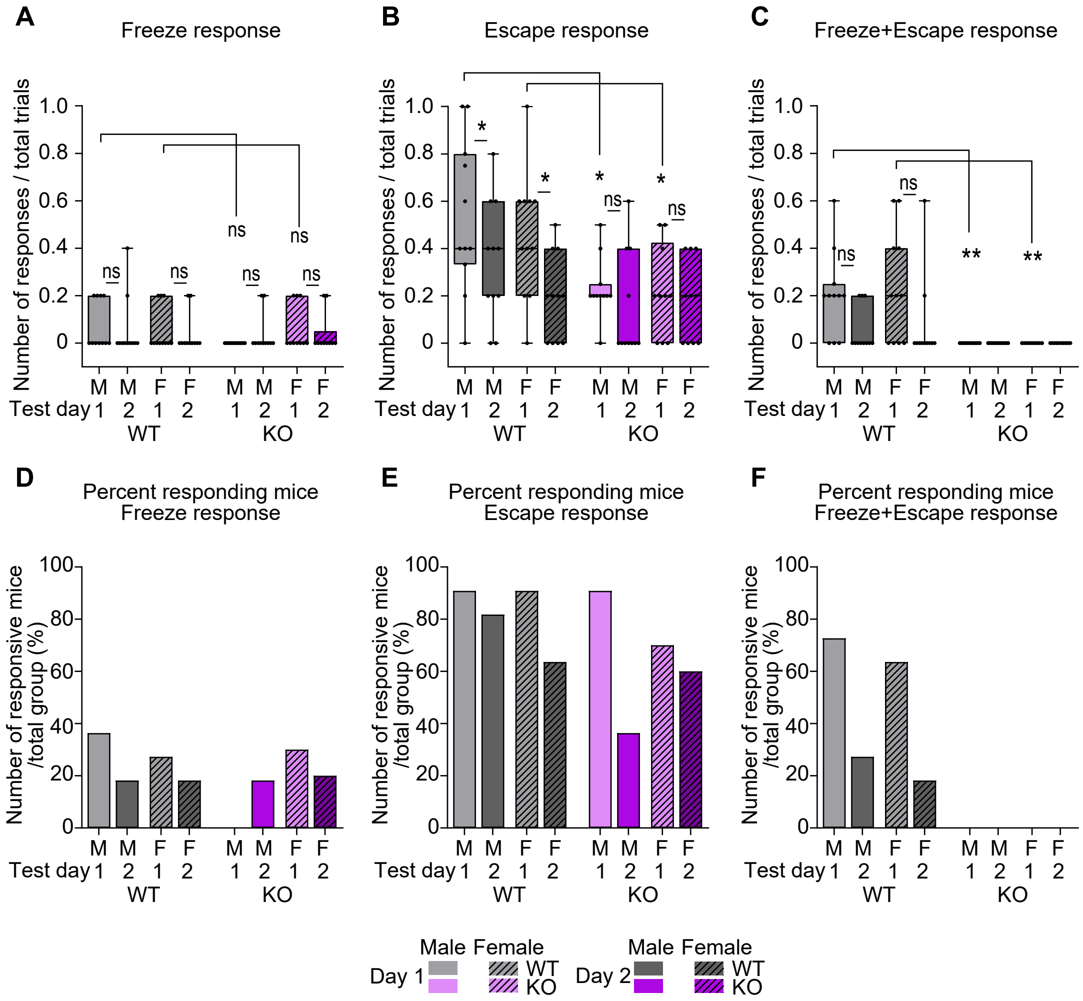

A
Freeze response
B
Escape response
C
Freeze+Escape response
Number of responses / total trials
ns
*
**
Test day
M 1 M 2 F 1 F 2
WT
KO
D
Percent responding mice
Freeze response
E
Percent responding mice
Escape response
F
Percent responding mice
Freeze+Escape response
Number of responsive mice /total group (%)
Male Female
Day 1
Day 2
WT
KO

**Figure S5. Visually evoked defensive behavior is disrupted in IP3R2 KO mice. A-C.** Average defensive responses separately quantified by response type as labeled (Freeze, Escape, and Freeze+Escape) across genotypes, sexes, and experimental days. Escape responses are significantly reduced and Freeze+Escape responses are abrogated in KO mice of both sexes. Freeze responses are not different between genotypes. Graphs show box with range, line is median. Circles on each box are responses of individual mice. **D-F.** Number of responders represented as percentage of mice responding to at least one trial per day out of total group number for each of the response types as labeled. All groups had similar percentage of responders with the exception of Freeze+Escape response where no KO mice were observed. Number of mice (N): 8-10/ per sex/ genotype. $^{*}P \leq 0.05$, $^{**}P<0.01$, $^{***}P<0.001$ by Mann-Whitney test. ns denotes non-significant results ($P>0.05$).

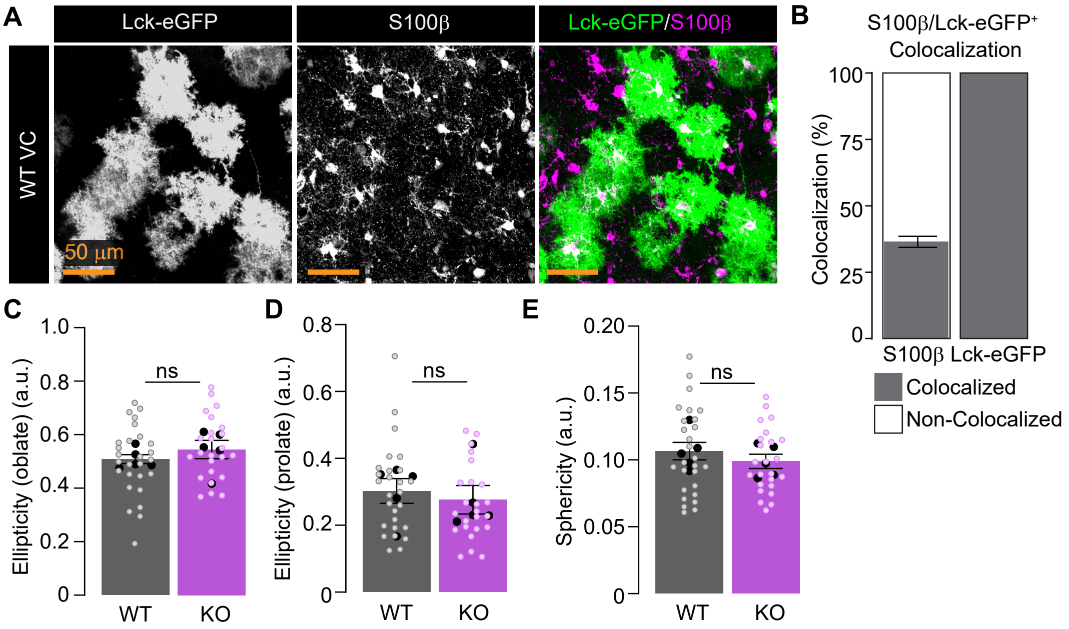
A
Lck-eGFP
S100β
Lck-eGFP/S100β
WT VC
50 μm
B
S100β/Lck-eGFP+
Colocalization
Colocalization (%)
100
75
50
25
0
S100β Lck-eGFP
Colocalized
Non-Colocalized
C
Ellipticity (oblate) (a.u.)
1.0
0.8
0.6
0.4
0.2
0
ns
WT
KO
D
Ellipticity (prolate) (a.u.)
0.8
0.6
0.4
0.2
0
ns
WT
KO
E
Sphericity (a.u.)
0.20
0.15
0.10
0.05
0
ns
WT
KO

**Figure S6. Reduced morphology in IP3R2 KO astrocytes. A-B.** Validation of the approach showing astrocyte specificity and sparsity of Lck-eGFP expression. Example images showing Lck-eGFP (green) and astrocyte specific marker S100β (magenta) (**A**) and quantification (**B**). Plot shows percentage of double labeled cells out of total astrocytes (S100β positive) or Lck-eGFP positive showing full overlap of Lck-eGFP expressing cells with S100β. **C-E.** Astrocytic shape characteristics including ellipticity and sphericity are unaltered by IP3R2 KO. Graphs show mean ± s.e.m. Black circles above each bar are average of signal in each mouse, colored open circles are data for each astrocyte. Number of mice/ group (N) N=5, number of astrocytes = 25-27. Scale bar = 50 μm. ns denotes non-significant results (P>0.05).
